# No evidence of genome-wide selection against sense transcription starts in human introns

**DOI:** 10.64898/2026.11.24.768857

**Authors:** Michał Wojciech Szcześniak, Elżbieta Wanowska

## Abstract

RNA polymerase II often starts transcription inside genes, usually at elements that initiate in both directions. An intragenic start pointing in the gene’s direction (a sense start) makes RNA that passes the gene’s splice sites and polyadenylation signals and may interfere with the gene. Introns may therefore have lost sequences that start sense transcription. In PRO-cap data from K562 and HCT116 cells, the sense start at intronic elements, measured against the antisense start, is a fifth to a third weaker than at intergenic elements of similar read depth. Sequence models that have never seen an element’s chromosome predict 30–42% of this bias from local DNA, except models trained on K562 applied to HCT116. However, introns as a whole show little or no depletion of predicted sense starts, and among 76,215 human genomes, variants predicted to create sense starts are not kept rarer than those creating antisense starts. In both cell lines, and in K562 GRO-cap data, strong measured start sites are rarer on the gene strand, mainly near the 5′ end of active genes. Thus, weaker sense starts appear to be a feature of individual active elements, partly predictable from their sequence, and we find no sign that selection has removed such sequences from introns.

## Introduction

Transcription by RNA polymerase II does not start only at the promoters of genes. Many sites inside genes, including enhancers in introns, also start transcription, and they usually do so in both directions, from two start sites a few hundred nucleotides apart (Seila et al. 2008; Kowalczyk et al. 2012; Core et al. 2014). Each direction is driven by its own core promoter, so the two starts of such an element are not simply mirror images, and their relative strength can differ (Core et al. 2014; Duttke et al. 2015). Cells limit transcription that starts inside genes. In yeast, methylation of histone H3 by Set2 during elongation recruits a deacetylase complex that keeps the chromatin inside genes closed (Carrozza et al. 2005), and in mouse embryonic stem cells the corresponding mark, H3K36me3, recruits DNA methylation that prevents spurious initiation inside genes (Neri et al. 2017). In human cells, depleting SETD2, the enzyme that deposits H3K36me3, lets transcription start inside at least 11% of active genes (Carvalho et al. 2013).

The direction of a start inside a gene matters. A start that points in the same direction as the gene, which we call a sense start, makes RNA that passes the gene’s downstream splice sites and polyadenylation signals. The 5′ splice sites recruit U1 snRNP, which protects RNA from early polyadenylation (Almada et al. 2013), so this RNA can be processed like the gene’s own messenger RNA, and its transcription can interfere with the gene; indeed, enhancers inside genes can lower the expression of their host genes (Cinghu et al. 2017). In contrast, a start that points in the opposite direction, an antisense start, usually meets an early polyadenylation signal, and its RNA is quickly degraded in the nucleus by the exosome (Preker et al. 2008; Almada et al. 2013; Ntini et al. 2013). Sense starts inside genes may therefore cost the cell more than antisense starts. If so, natural selection could remove from introns the sequences that are able to start sense transcription. Selection of this kind has been reported in bacteria and archaea, whose genomes contain fewer short sequences resembling promoters or transcription factor binding sites than expected by chance (Hahn et al. 2003; Froula and Francino 2007). Such selection would need to act all the time, because in bacteria random sequences can rapidly evolve into promoters (Yona et al. 2018).

A difference between the two strands of introns could also arise without selection. Transcription changes how DNA mutates: over evolutionary time, the coding strand of genes gains an excess of G and T over A and C, a pattern known as keto skew (Green et al. 2003; Polak and Arndt 2008), and the size of this skew follows how strongly a gene is transcribed in the germline (McVicker and Green 2010). If DNA of this composition happens to start transcription less well, the gene strand of introns would start less sense transcription for purely mutational reasons. Finally, the weaker sense start may be a property of the elements themselves, set by the sequence of their two core promoters (one for each direction) or by their surroundings in the cell, such as the histone marks and DNA methylation that transcription of the host gene leaves behind, without any genome-wide change in intronic DNA.

In previous work (MW Szcześniak and E Wanowska, unpubl.), we used PRO-cap data, which map the capped 5′ ends of newly made RNA and so show where transcription starts (Kwak et al. 2013), and observed that, in start-site pairs inside introns (the two divergent starts of one element), the sense start is weaker than the antisense start. Here we ask which of these explanations accounts for this bias, using three kinds of evidence. First, two deep-learning models that predict transcription start sites from DNA sequence, Puffin-D (Dudnyk et al. 2024) and ProCapNet (Cochran et al. 2024), show how much of the bias can be predicted from the sequence of the elements, and whether introns as a whole contain fewer sequences able to start sense transcription. Second, start sites measured in the same cells show which of the model predictions hold in real data. Third, genetic variation in 76,215 human genomes (Guez et al. 2026) shows whether new sense starts are removed from the population by selection.

We find that the bias is robust, and that sequence models predict about a third to two fifths of it from the local sequence of the elements; models trained on K562 cells, however, predicted little of it in HCT116 cells. In contrast, we find no evidence of genome-wide selection: introns as a whole show little or no depletion of sequences able to start sense transcription, and variants that create sense starts are not kept rare in the population. In both cell lines, strong start sites measured by PRO-cap are nevertheless less frequent on the gene strand, mainly near the 5′ end of active genes, and a second assay, GRO-cap, confirms this in K562. Together, these results suggest that the weaker sense start is a feature of the individual elements that start transcription inside genes in these cells, partly predictable from their sequence and largest near the start of the host gene, rather than the result of selection that removed sequences able to start sense transcription from the genome.

## Results

### Sense starts at intronic elements are weaker than at intergenic elements

We used start-site pairs identified in PRO-cap data from two human cell lines, K562 and HCT116. In each pair, two starts a few hundred nucleotides apart point away from each other (Fig. 1A). In a pair inside an intron of a protein-coding gene, the sense start points in the same direction as the gene, and the antisense start points in the opposite direction. For each pair, we measured the asymmetry as the log2 ratio of PRO-cap reads at the sense start to reads at the antisense start. Because the two starts share their surroundings, this ratio cancels effects that act on both starts equally, such as local chromatin and overall read depth. As the reference, we used intergenic pairs, which lie more than 5 kb from any protein-coding gene; in these pairs neither direction is special, so their asymmetry should be centered on zero. We express the bias as the percentage by which the ratio of sense to antisense reads is lower in intronic pairs than in intergenic pairs. We analyzed 1,863 intronic pairs in K562 and 2,858 in HCT116, against 3,933 and 6,193 intergenic pairs. The main comparisons use the intronic pairs in active host genes (the genes that contain the pairs), that is, genes with at least five PRO-cap reads at their annotated start (Methods): 1,611 pairs in K562 and 2,489 in HCT116.

**Figure 1.**
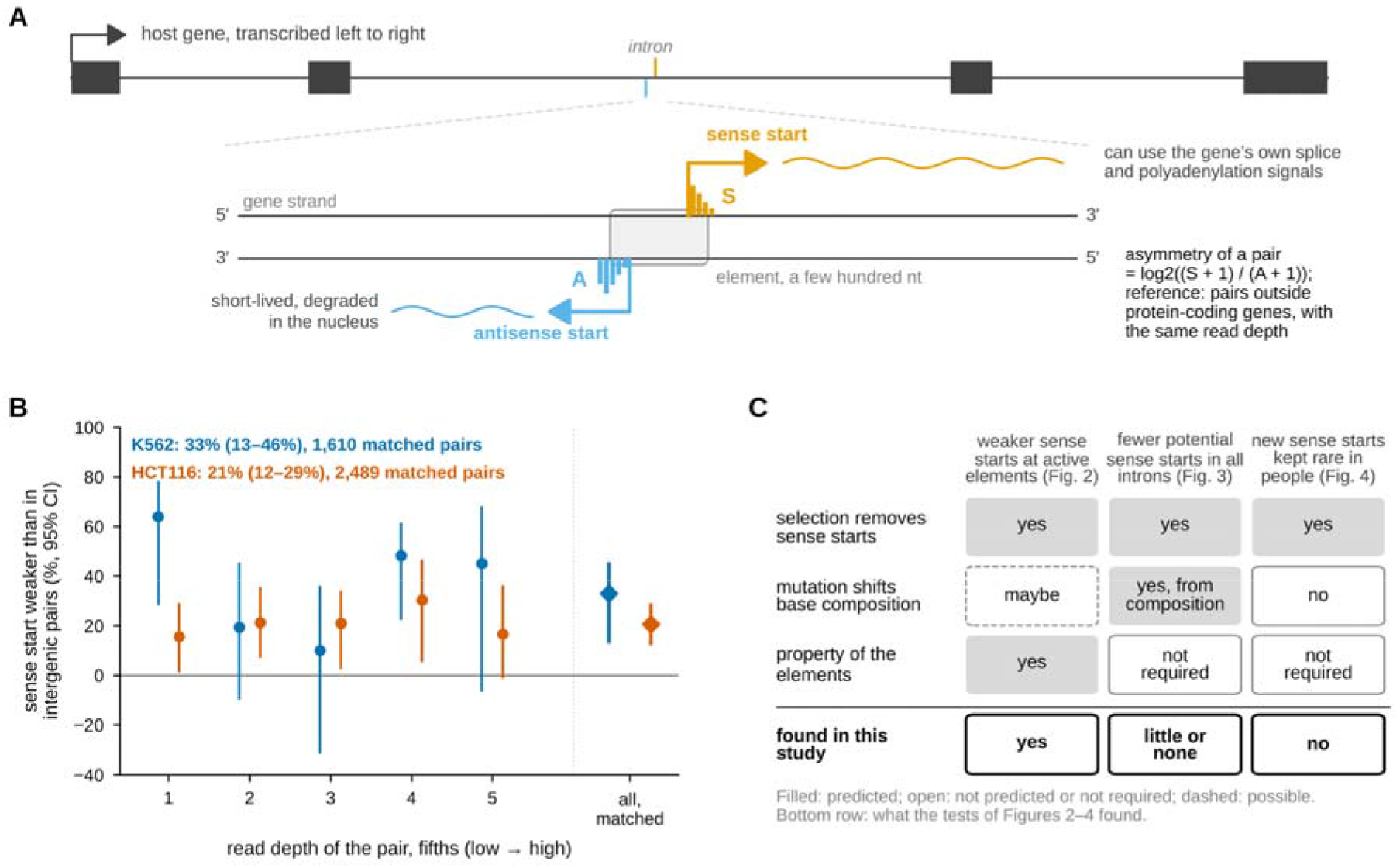
Sense starts at intronic elements are weaker than at intergenic elements. (A) A pair of transcription starts inside an intron. The sense start (orange) points in the same direction as the host gene, so its RNA can reach the gene’s downstream splice sites and polyadenylation signals; the antisense start (blue) makes RNA that is usually degraded in the nucleus. The asymmetry of a pair is the log2 ratio of PRO-cap reads at its sense (S) and antisense (A) start. (B) Percentage by which the ratio of sense to antisense reads is lower in intronic pairs in active host genes than in intergenic pairs, within five bands of read depth (circles) and after one-to-one depth matching (diamonds); K562, blue; HCT116, orange-red (these colors are unrelated to panel A). Values are 100 × (1 − 2), where d is the difference in median log2 ratio between intronic and intergenic pairs; bars, 95% CI from 10,000 resamples of host genes (intergenic pairs within bands: 1-Mb blocks), converted to percent in the same way; the axis stops at 100%, the largest possible value. Pairs per band: K562, 157–457 intronic and 664–952 intergenic; HCT116, 347–604 and 1,135–1,388. Matched comparison: n = 1,610 (K562) and 2,489 (HCT116) intronic pairs, each with one intergenic pair. (C) Three explanations of the bias (rows) and what each predicts for the three tests of this study (columns; Figs. 2–4); ‘not required’, the explanation does not need this result. The bottom row gives what the tests found.

Pairs with few reads give noisy ratios, and intronic pairs carry fewer reads than intergenic pairs. We therefore compared only pairs of similar read depth, in two ways. First, we divided the pairs into five bands of read depth. In every band, in both cell lines, the sense start of intronic pairs was weaker (by 10–64% in K562 and 16–30% in HCT116, depending on the band), although in four of the ten bands the 95% confidence interval (CI) included zero (Fig. 1B). Second, we matched each intronic pair to one intergenic pair of nearly the same read depth (within 10%); all but one of the 1,611 K562 pairs found such a partner. After matching, the sense start was weaker by a third in K562 (33%, 95% CI 13–46%) and by a fifth in HCT116 (21%, 12–29%; two- sided Wilcoxon signed-rank test on the matched differences, P = 0.001 in K562 and P = 2 × 10 in HCT116). Because pairs in the same host gene may behave alike (in K562, the 1,610 matched pairs lie in 964 host genes), we calculated the CIs by resampling whole host genes, each with all of its pairs. Resampling single pairs gave almost the same intervals (Supplemental Table S1).

Most pairs are not alternative promoters of their host genes: only 6–8% had a start within 100 nt of an annotated transcript start, and removing such pairs, or pairs within 1 kb, left the bias almost unchanged (Supplemental Results; Supplemental Table S1).

Three explanations of this bias make different predictions (Fig. 1C), and more than one of them may contribute. First, if selection removes sequences that start sense transcription, active elements should have weaker sense starts, introns as a whole should contain fewer potential sense starts, and variants that create new sense starts should be kept rare in the population. Second, if the cause is the mutational shift in base composition of the gene strand (the strand that runs in the direction of the gene), any DNA of the same composition should show the bias, without selection. Third, if the cause lies in the elements themselves, for example in a sense core promoter weaker than the antisense one, active elements should have weaker sense starts, but introns as a whole need not contain fewer potential sense starts, and selection is not needed. The next sections test these predictions in turn (bottom row of Fig. 1C).

### The weaker sense start does not detectably depend on host activity and is partly predicted from the element’s sequence

We next asked whether the bias depends on transcription of the host gene. Marks that transcription leaves inside genes suppress initiation there in yeast (Carrozza et al. 2005), in mouse embryonic stem cells (Neri et al. 2017) and in human cells (Carvalho et al. 2013). Whether these marks act more strongly on sense than on antisense starts is not known. If they do, the bias should grow with the activity of the host gene and should disappear where the host is silent. We therefore repeated the depth-matched comparison in groups of host activity, measured as PRO-cap reads at the host gene’s own start. The point estimate showed a weaker sense start in every group, and the bias did not change detectably with host activity (Fig. 2A; Spearman correlation between host activity and the matched difference in asymmetry, ρ = 0.00, 95% CI −0.04 to 0.04, in K562 and ρ = −0.02, −0.05 to 0.02, in HCT116). In silent hosts, the bias was at least as large as in active hosts (47% compared with 33% in K562, and 24% compared with 21% in HCT116), although the few pairs in silent hosts made this difference uncertain (Supplemental Table S1). The bias also did not differ detectably between pairs with and without nearby H3K36me3, a chromatin mark of transcription elongation (Supplemental Table S1). Our data therefore give no evidence that transcription of the host gene imposes the bias, although they cannot exclude a modest effect; an earlier analysis of the same pairs without depth matching reached the same conclusion (MW Szcześniak and E Wanowska, unpubl.).

**Figure 2.**
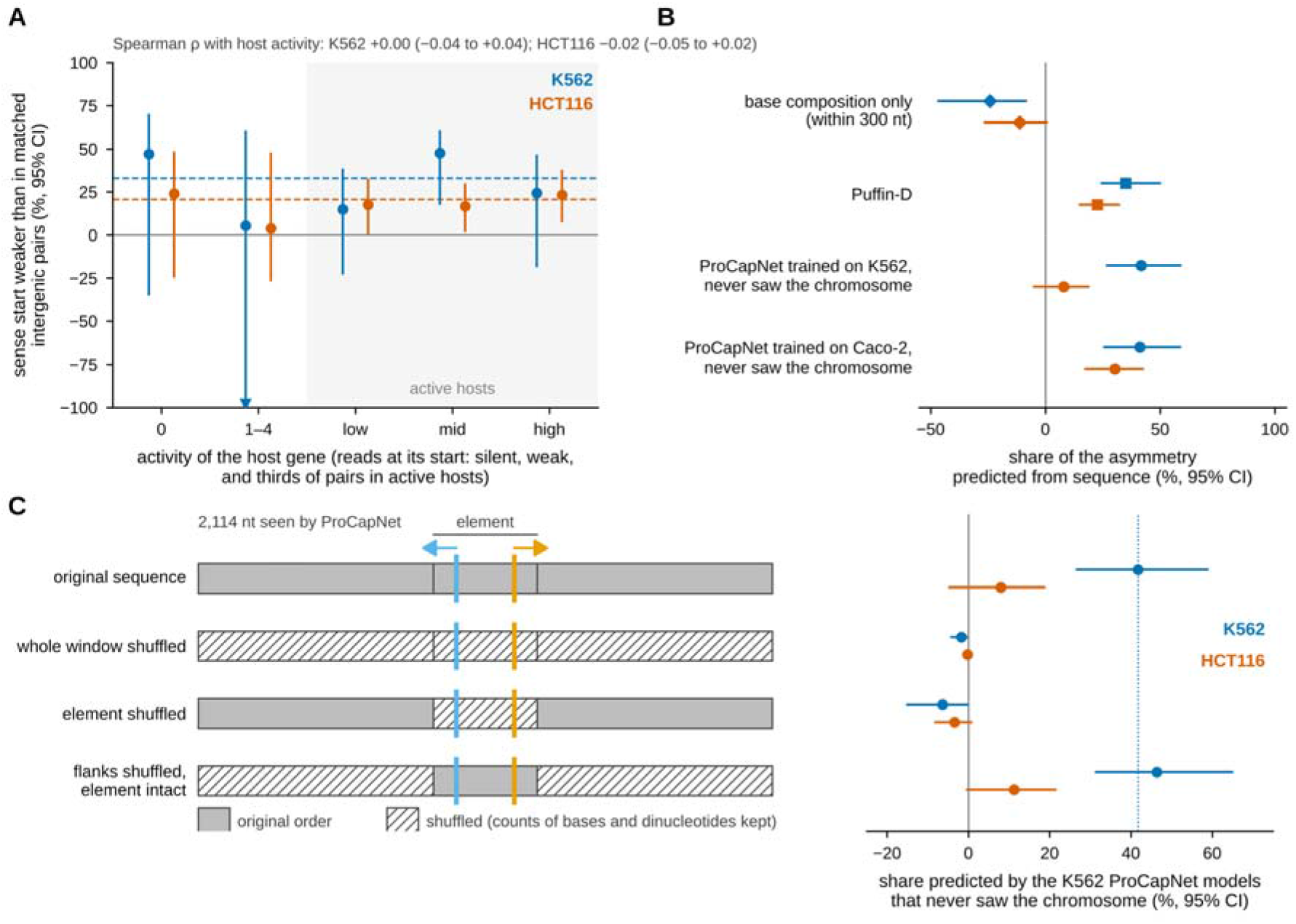
The weaker sense start does not detectably depend on host activity and is partly predicted from the element’s own sequence. K562, blue; HCT116, orange-red. (A) Percentage by which the ratio of sense to antisense reads is lower in intronic pairs than in depth-matched intergenic pairs, by activity of the host gene: silent (no PRO-cap reads at the host’s start), weakly active (1–4 reads) and thirds of host activity among pairs in active hosts. Points, median-based percentages as in Figure 1B; bars, 95% CI from 10,000 resamples of host genes; dashed line, all intronic pairs in active hosts; triangle, interval continues below the axis. Above the panel, Spearman correlation between host activity and the difference in asymmetry between each intronic pair and its matched intergenic pair, with 95% CI. Intronic pairs per group, each matched to one intergenic pair: K562, 148 (silent), 104 (weakly active) and 536–537 (each third); HCT116, 214, 155 and 829–830. (B) Share of the measured asymmetry of intronic pairs in active hosts that is predicted from sequence: by base composition alone (excess of T over A and of G over C within 300 nt, in a linear model; negative values mean that this model predicted a stronger, not a weaker, sense start), by Puffin-D, and by ProCapNet models trained on K562 or on Caco-2 (colorectal) PRO-cap that never saw the pair’s chromosome. Puffin-D was trained on all chromosomes except 8–10, which hold 16% (K562) and 14% (HCT116) of the pairs. Share = calibrated mean predicted asymmetry divided by mean measured asymmetry (calibration on intergenic pairs, each counted in both orientations; Methods). Bars, 95% CI from 2,000 resamples of host genes, with intergenic pairs resampled in 1-Mb blocks (composition: 1,000 resamples of host genes and of intergenic pairs). n = 1,611 (K562) and 2,489 (HCT116) pairs. (C) Where in the sequence the prediction lies. Left, the 2,114 nt seen by ProCapNet around a pair (schematic), with the shuffled part hatched; orange and blue arrows, the sense and antisense start (colors unrelated to the cell lines); shuffles keep the count of every base and of every dinucleotide, and runs of N stay in place; element, from 100 nt before the first start to 100 nt after the second. Right, share of the asymmetry predicted after each shuffle by the K562 models that never saw the chromosome (five shuffles per pair, predicted signal averaged; calibration, CI and n as in B). Dotted line, share for the original sequence in K562.

We then asked how much of the bias can be predicted from the DNA sequence of the elements. We used two deep-learning models that predict, from DNA sequence alone, where transcription starts and how strongly; an asymmetry that a model predicts therefore comes from the DNA that the model sees. The two models see different amounts of DNA: Puffin-D sees 100 kb around each position (Dudnyk et al. 2024), and ProCapNet sees 2.1 kb (Cochran et al. 2024). ProCapNet was trained on K562 PRO-cap from the same ENCODE experiment that gave our K562 pairs (Methods). For each pair, we predicted the strength of both starts and converted the predicted asymmetry to the scale of the measured asymmetry, using intergenic pairs as the reference (calibration; Methods). The share of the bias predicted from sequence is the converted prediction for intronic pairs divided by their measured asymmetry (100%, the whole bias; a negative share, the opposite direction).

Because a model may partly memorize the start sites on which it was trained, we used held-out ProCapNet models, which had never seen the chromosome of the pair (ProCapNet was trained seven times, each time without a different group of chromosomes). These held-out models reproduced about two fifths of the bias in K562 (42%, 95% CI 26–59%; Fig. 2B). Models that had seen the chromosome of the pair reproduced more (63%, 46–83%; Supplemental Fig. S1B), as expected if they partly memorize their training data. Puffin-D reproduced 35% (24– 50%) of the bias in K562 and 23% (15–33%) in HCT116. These two values may be inflated in the same way, because 84% of the matched pairs in K562 and 86% in HCT116 lie on chromosomes on which Puffin-D was trained (all chromosomes except 8, 9 and 10; Methods). Part of the Puffin-D share may also come from the host gene, whose promoter often lies within the 100 kb that Puffin-D sees (see below). Two other ways of measuring the share, a rank- based comparison that needs no calibration and a calibration within bands of read depth, led to the same conclusion (Supplemental Fig. S1B).

The share predicted by ProCapNet depended on the cells on which the model was trained. In HCT116, the ProCapNet models trained on K562 reproduced little of the bias (8%, 95% CI −5 to 19%; Fig. 2B). ENCODE also provides ProCapNet models trained, with the same groups of chromosomes held out, on PRO-cap from Caco-2, a colorectal cancer cell line like HCT116. Held-out Caco-2 models predicted about a third of the bias in HCT116 (30%, 17–43%) and as much as the K562 models in K562 (41%, 25–59%). Because the Caco-2 models were trained on other cells in another experiment, these two estimates share no data between training and testing.

Does the order of the bases matter? We shuffled the DNA that ProCapNet sees around each pair, keeping how often each base and each two-base combination (such as CG) occurs, and scored the shuffled sequences with the held-out models (Fig. 2C). Shuffling the whole 2.1-kb window, or only the element, removed the predicted bias, so composition alone cannot produce it. Both shuffles, however, also destroy the core promoters of the two starts. When only the flanks were shuffled, that is, everything except the element (from 100 nt before the first start to 100 nt after the second), ProCapNet still predicted 46% of the bias in K562, compared with 42% for the original sequence. The information that ProCapNet uses thus lies within the element itself, within a few hundred nucleotides of the two starts. In HCT116, the K562 models predicted too little of the bias (8%) for this test.

Three alternatives did not explain the bias (Supplemental Results). Base composition near the pairs predicted a slightly stronger, not a weaker, sense start (share −24% in K562 and −11% in HCT116; Fig. 2B); known core promoter elements, such as the TATA box and the initiator, were not more common at the antisense start than at the sense start (Supplemental Fig. S2); and removing pairs that overlap transposable elements left the bias almost unchanged (31% in K562 and 25% in HCT116). A substantial part of the weaker sense start can therefore be predicted from the sequence of the active elements. If the bias reflected a removal of sense-start sequences from introns in general, however, introns as a whole should also contain fewer potential sense starts. We tested this next.

### Introns as a whole show little or no depletion of potential sense starts

If selection removes sequences that start sense transcription, the effect should not be limited to the few thousand elements that are active in one cell line: introns as a whole should contain fewer positions from which sense transcription could start. We therefore scored every position of the 22 autosomes, on both strands, with both models, and counted potential start sites; we call this genome-wide count the census. A position was a potential start site when its predicted signal reached the highest 0.1%, 0.03% or 0.01% of predictions on chromosomes 8 and 9 (weak, medium and strong sites; Methods). We counted these sites on the gene strand and on the opposite strand of the introns of 14,267 protein-coding genes, leaving out exons, annotated transcript starts and regions where another gene overlaps. A potential start site on the gene strand is a potential sense start. As a control, we counted sites in the same way on the two strands of DNA more than 10 kb from any annotated gene, where the two strands should not differ. The two models differ in how much DNA they see, 2.1 kb and 100 kb (Fig. 3A). We took the Puffin-D predictions for each position from the central 50 kb of a 100-kb window, so the window extended at least 25 kb and at most 75 kb to each side of the position, the two sides together making 100 kb. Puffin-D therefore saw the promoter of the host gene for every position within 25 kb of that promoter, and for some positions up to 75 kb away.

**Figure 3.**
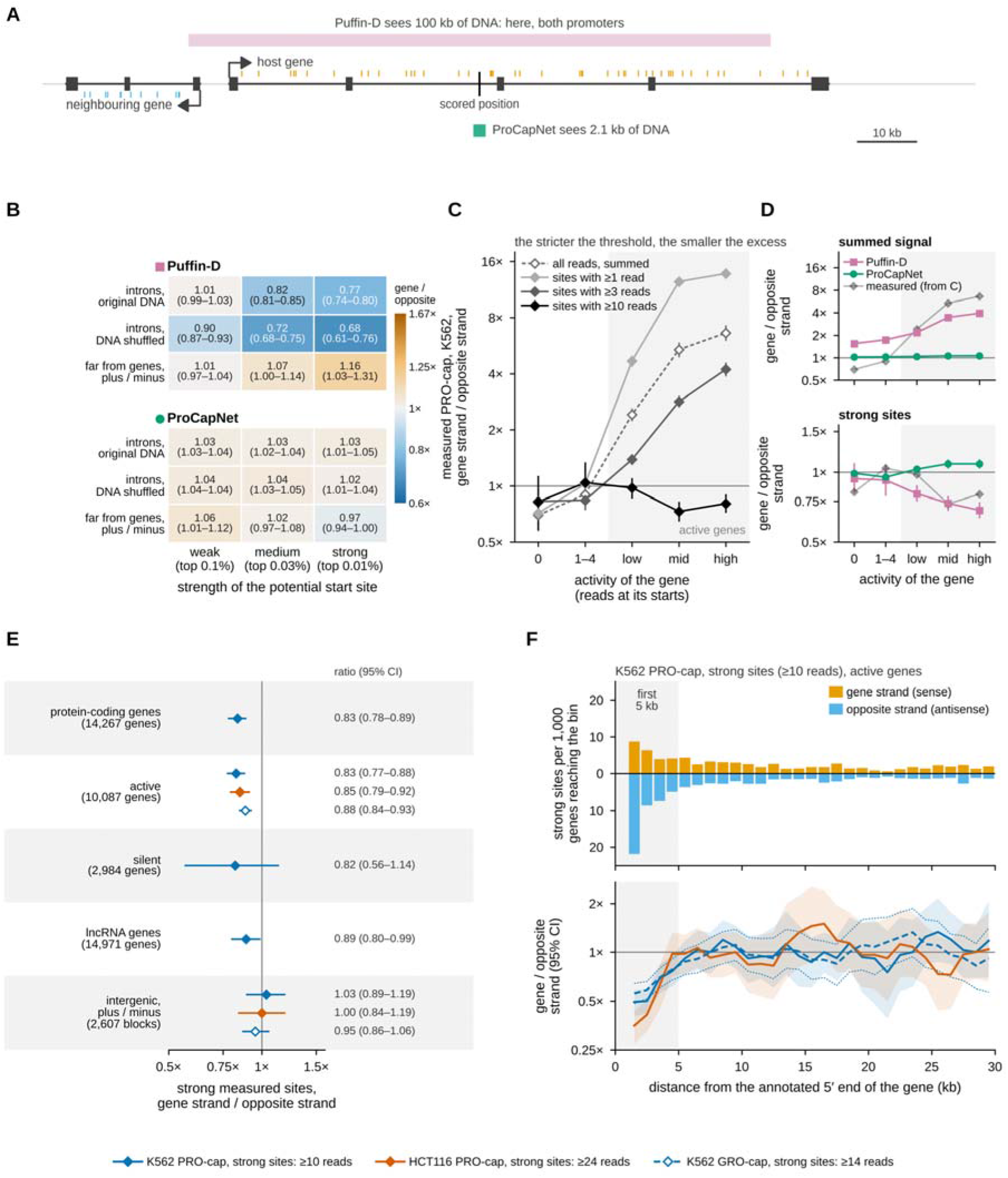
Potential and measured start sites on the two strands of introns. (A) How much DNA each model sees when it scores one intronic position, drawn to scale: Puffin-D, 100 kb, which here includes the promoters of the host gene and of its neighbor; ProCapNet, 2.1 kb. Pink bar, the DNA seen by Puffin-D; green box, the DNA seen by ProCapNet. Ticks, PRO-cap reads along the transcribed genes: orange above the line, on the host gene’s strand; blue below the line, on the opposite strand, from the neighboring gene (illustration; the models see only DNA). (B) Potential start sites of Puffin-D and ProCapNet, gene strand divided by opposite strand, for weak, medium and strong sites (highest 0.1%, 0.03% and 0.01% of predictions on chromosomes 8 and 9): in introns of 14,267 protein-coding genes (Puffin-D, 14,266) with their original DNA; in the same introns after shuffling their DNA with dinucleotide composition preserved; and, as a control expected to be 1, in DNA far from genes (plus strand divided by minus strand; 2,606 1-Mb blocks for Puffin-D and 2,607 for ProCapNet). Each cell gives the ratio and its 95% CI; color, the same ratio on a logarithmic scale (blue, fewer sites on the gene strand; orange, more). (C) Measured PRO-cap signal in K562, gene strand divided by opposite strand, in introns grouped by the activity of the gene (reads at its annotated starts: silent, 0; weakly active, 1–4; and thirds of active genes; gray shading, active genes): all reads summed (dashed line, open diamonds) and sites with at least 1, 3 or 10 reads (light gray to black). (D) The same ratio predicted by Puffin-D (squares) and ProCapNet (circles) in the same groups of genes, for the summed signal (top) and for strong potential sites (highest 0.01%; bottom), next to the measured curves of C (faint: all reads summed, top; sites with at least 10 reads, bottom). (E) Strong measured sites, gene strand divided by opposite strand, in sets of genes and in intergenic DNA (plus strand divided by minus strand; number of genes or of 1-Mb blocks in brackets) in K562 PRO-cap and, for active protein-coding genes and intergenic DNA, in HCT116 PRO-cap (10,538 genes) and K562 GRO-cap (9,907 genes); at right, the ratio and its 95% CI. The comparisons in active genes in HCT116 and in K562 GRO-cap were recorded before the data were seen. (F) Strong measured sites in active protein-coding genes by distance from the annotated 5′ end of the gene, in 1-kb bins up to 30 kb (eligible positions lie at least 1 kb from annotated transcript starts; sites farther than 30 kb, about half of all, are included in Supplemental Fig. S5B). Top, K562 PRO-cap: sites per 1,000 active genes whose annotated span reaches the bin, on the gene strand (above the line) and on the opposite strand (below). Bottom, gene strand divided by opposite strand in 3-kb windows centered on each bin (the first window spans 1–3 kb), with its 95% CI (shading; dotted lines for K562 GRO-cap), in the three data sets of E; gray shading, the first 5 kb. Panel F is descriptive and was drawn after the analyses of distance (Methods). Strong measured sites: at least 10 reads in K562 PRO-cap, 24 in HCT116 PRO-cap and 14 in K562 GRO-cap. All strand ratios on a logarithmic axis; values below 1 mean fewer sites on the gene strand. 95% CI from 2,000 bootstrap resamples of genes (in F, within thirds of gene activity; intergenic DNA, 1-Mb blocks). Genes per group in C and D: 1,196–3,366 (silent genes: 2,983, because one gene is missing from the Puffin-D tables; 2,984 in E). Numbers of sites in each group are given in the Source Data.

The two models gave different answers (Fig. 3B). Puffin-D predicted about a fifth fewer medium and strong sites on the gene strand than on the opposite strand (18% and 23% fewer). ProCapNet predicted no such deficit: at every threshold, the gene strand carried slightly more sites than the opposite strand (about 3% more). To separate the effect of particular sequences from that of base composition (the gene strand of introns is richer in G and T), we shuffled the DNA of every intron, keeping its dinucleotide composition, and scored it again: if introns had lost sequences able to start sense transcription, real introns should carry fewer gene-strand sites than their shuffled versions. According to ProCapNet, real introns carried only slightly fewer weak and medium sites than shuffled introns (the excess of sites on the gene strand, about 3%, was 0.8 and 1.5 percentage points smaller than in shuffled introns) and no fewer strong sites; outside repeat-derived sequence, real and shuffled introns did not differ (Supplemental Table S1). For Puffin-D, the deficit was larger in shuffled introns than in real ones, so it cannot come from particular intronic sequences. It must come from something that shuffling keeps, such as base composition or the DNA outside the intron (see below).

How large a strand difference can arise where none is expected? In the intergenic control, where neither strand is special, the ratio of sites on the plus strand to sites on the minus strand ranged from 0.97 to 1.16, and some of the CIs excluded 1 (Fig. 3B). The strand differences that ProCapNet predicted in introns lie within this range. According to the model that sees only local sequence, therefore, introns show little or no depletion of sequences able to start sense transcription.

Why did Puffin-D give a different answer? Summed over all intronic positions, Puffin-D predicted almost three times more signal on the gene strand than on the opposite strand (2.8×), whereas ProCapNet predicted almost equal amounts (1.04×). Start sites measured by PRO-cap in K562 at the same positions (data from the experiment that gave the K562 pairs) showed a large strand difference too: in active genes, sites with at least one read were about ten times more frequent on the gene strand than on the opposite strand, and this excess grew with the activity of the gene (Fig. 3C). Besides discrete start sites, PRO-cap therefore records a diffuse signal along the gene strand that grows with transcription of the gene; we call it the transcription signal. This signal may be background from uncapped RNA made by polymerases elongating through the gene, which the selection for capped 5′ ends does not remove completely, or genuine low-level initiation coupled to elongation; our data do not separate the two. The transcription signal fades at higher read thresholds (Fig. 3C). The summed signal predicted by Puffin-D also rose with the activity of the gene, although the model sees only DNA, whereas the signal predicted by ProCapNet stayed nearly equal on the two strands (Fig. 3D). Because the Puffin-D window often includes the promoter of the host gene, the model has apparently learned from the DNA where genes are transcribed; inside genes, Puffin-D therefore mixes the transcription signal with the ability of the sequence to start transcription and cannot give a simple count of potential starts.

The same signal explains why weak measured sites were rarer on the gene strand of silent genes: the deficit followed the strand reached by transcription from active neighboring genes (Supplemental Results; Supplemental Fig. S3).

### Strong measured start sites are less frequent on the gene strand in both cell lines, most clearly near the start of active genes

Strong measured start sites, those with at least ten reads, carried much less of the transcription signal (Fig. 3C) and therefore allow a cleaner comparison of the two strands. In introns of protein-coding genes in K562, the gene strand carried about a sixth fewer strong sites than the opposite strand (16.5%, 95% CI 10.8–22.3%). Introns of lncRNA genes showed a smaller deficit (11%, 1–20%), and intergenic DNA showed no detectable difference (Fig. 3E). The deficit was reproduced in HCT116, in an analysis recorded before seeing the data (Methods; strong sites, at least 24 reads): in active protein-coding genes, the gene strand carried 15% fewer strong sites than the opposite strand (95% CI 8–21%; 927 and 1,092 sites), close to the 17% (12–23%) in active genes of K562, whereas intergenic DNA again showed no difference (0%, −19 to 16%; Fig. 3E).

The deficit was not spread evenly over genes (Fig. 3C). It was clear in the middle and the most active third of genes (27%, 95% CI 18–35%, and 20%, 10–28%, respectively), but not detectable in the least active third (2%, −10 to 14%) or in weakly active genes (1–4 reads at their starts; −4%, −34 to 20%). In silent genes, the estimate (18%, −13 to 42%) was close to that in all active genes taken together (17%, 95% CI 12–23%; Fig. 3E), but imprecise. In HCT116, the deficit likewise grew with activity, from 3% (−10 to 15%) in the least active third to 13% (2–24%) and 29% (17–38%) in the middle and the most active third. Which model reproduced this pattern (Fig. 3D)? Puffin-D did, in part: the deficit of strong sites that Puffin-D predicted also rose with gene activity, from 6% in silent genes to 31% in the most active third.

ProCapNet, in contrast, predicted about 9% more strong sites on the gene strand in the middle and the most active third. In the census, Puffin-D’s knowledge of where genes are transcribed was a drawback. Here, the same knowledge is informative. Part of the deficit in the more active genes may therefore be linked to the activity of the host gene, which Puffin-D sees, rather than to the local sequence, which is all that ProCapNet sees.

DNA methylation inside genes, which can prevent spurious initiation (Neri et al. 2017), showed no clear link with the deficit: the planned comparison found no difference, and a comparison chosen afterwards found a moderate, uncertain one (Supplemental Results; Supplemental Fig. S4).

A second mark, H3K36me3, is laid down during elongation, helps suppress initiation inside transcribed genes (Carrozza et al. 2005; Carvalho et al. 2013) and rises toward the 3′ end of active genes (Bannister et al. 2005). If H3K36me3 suppressed sense starts, the deficit should be larger inside H3K36me3 peaks than outside them, and larger in the 3′ fifth of a gene than in the 5′ fifth. We recorded these two predictions, together with the replication in HCT116, before seeing the data and tested them in active protein-coding genes of both cell lines (Methods). The first test was not informative: only 104 strong sites in K562 and 25 in HCT116 lay inside peaks, and the difference inside minus outside peaks was very uncertain (K562, −18.1 percentage points, 95% CI −73.3 to 18.0; HCT116, 6.4, −90.3 to 51.0). The second test gave the opposite of the prediction. The deficit was large in the 5′ fifth of genes (35%, 95% CI 26–42%, in K562 and 37%, 28–45%, in HCT116) and absent in the 3′ fifth (5%, −16 to 22%, and −4%, −28 to 14%), so the difference 3′ minus 5′ was negative (−30.0 percentage points, −51.9 to −10.6, P = 0.002, and −41.0, −66.8 to −20.0, P = 0.001; two-sided bootstrap over genes; Supplemental Fig. S5A). The deficit of strong sense sites therefore lies toward the 5′ end of genes, whereas H3K36me3 rises toward the 3′ end.

Where along the gene does the deficit lie? In analyses chosen after seeing the results above, and therefore exploratory (Methods), we measured the distance of each strong site from the annotated 5′ end of the host gene. In both cell lines, the deficit was confined to the first 5 kb (46%, 95% CI 38–54%, in K562 and 56%, 47–65%, in HCT116) and was close to zero farther away (6%, −2 to 12%, and 3%, −6 to 10%; Supplemental Fig. S5B). In 1-kb bins, the deficit was largest within 1–3 kb of the 5′ end, where the gene strand carried about half as many strong sites as the opposite strand (ratio 0.49, 95% CI 0.40–0.59, in K562 and 0.35, 0.27–0.44, in HCT116), and it had faded by about 5 kb (Fig. 3F). In K562, both strands carried more strong sites near the 5′ end than farther along the gene, the opposite strand more so (Fig. 3F, top). The weaker sense start of the pairs was also strongest in the first 5 kb (65%, 44–79%, and 51%, 35– 66%) and smaller farther away (21%, −6 to 37%, and 14%, 4–24%; Supplemental Fig. S5C).

Most strong sites and pairs near the start of a gene lay within 2 kb of a CpG island, where antisense initiation is common (Lavender et al. 2016). The concentration in the first 5 kb, however, remained more than 2 kb from any island and did not grow detectably with gene activity, whereas islands farther from the start did not raise the strand difference (Supplemental Results; Supplemental Fig. S5D, E).

To check these findings with a second assay, we repeated the analysis of strong sites with K562 GRO-cap data (Core et al. 2014), produced with an earlier protocol and transferred from the hg19 to the hg38 genome (Methods). Before the analysis, we recorded three predictions, and all three held. In active protein-coding genes, the gene strand carried 12% fewer strong sites than the opposite strand (95% CI 7–16%; Fig. 3E); the deficit was smaller in the 3′ fifth of genes than in the 5′ fifth (−21.3 percentage points, −37.9 to −6.4, P = 0.004; Supplemental Fig. S5A); and the deficit was larger within the first 5 kb than farther away (35.1 percentage points, 26.1–43.8, P < 0.001; Fig. 3F; Supplemental Fig. S5B). Intergenic DNA again showed no difference (5%, −6 to 14%).

Whether the deficit of strong measured sites depends on base composition remains open. The deficit was larger in genes whose gene strand carries the largest excess of G and T, but the CI included zero, and the two models responded to this composition in opposite directions (Supplemental Fig. S6).

### No sign of selection against new sense start sites in human populations

The census found no clear depletion of potential sense starts, but it could miss selection that began too recently to have changed the sequence of introns, so we also looked for selection directly, in present-day human variation. The test rests on a simple idea: selection removes harmful variants before they become common, so a harmful class of variants contains more variants seen in only one person (singletons) than a harmless class. The proportion of singletons, however, also depends on how often a site mutates. A variant at a site that mutates often, such as a CpG, arises repeatedly in different people and is therefore less often a singleton, even without selection. We therefore compared each class of variants with variants that have the same mutation type (for example, C to T) and the same neighboring bases (three- base context) but no predicted effect. The proportion of singletons in a class minus the proportion in these matched variants is the excess of singletons (Lek et al. 2016; Fig. 4A); a positive excess suggests selection against the class. We used 215.5 million single-nucleotide variants from 76,215 genomes in gnomAD v4.1 (Methods) and classified each variant by the effect on initiation predicted for each strand, separately with ProCapNet and with Puffin-D.

**Figure 4.**
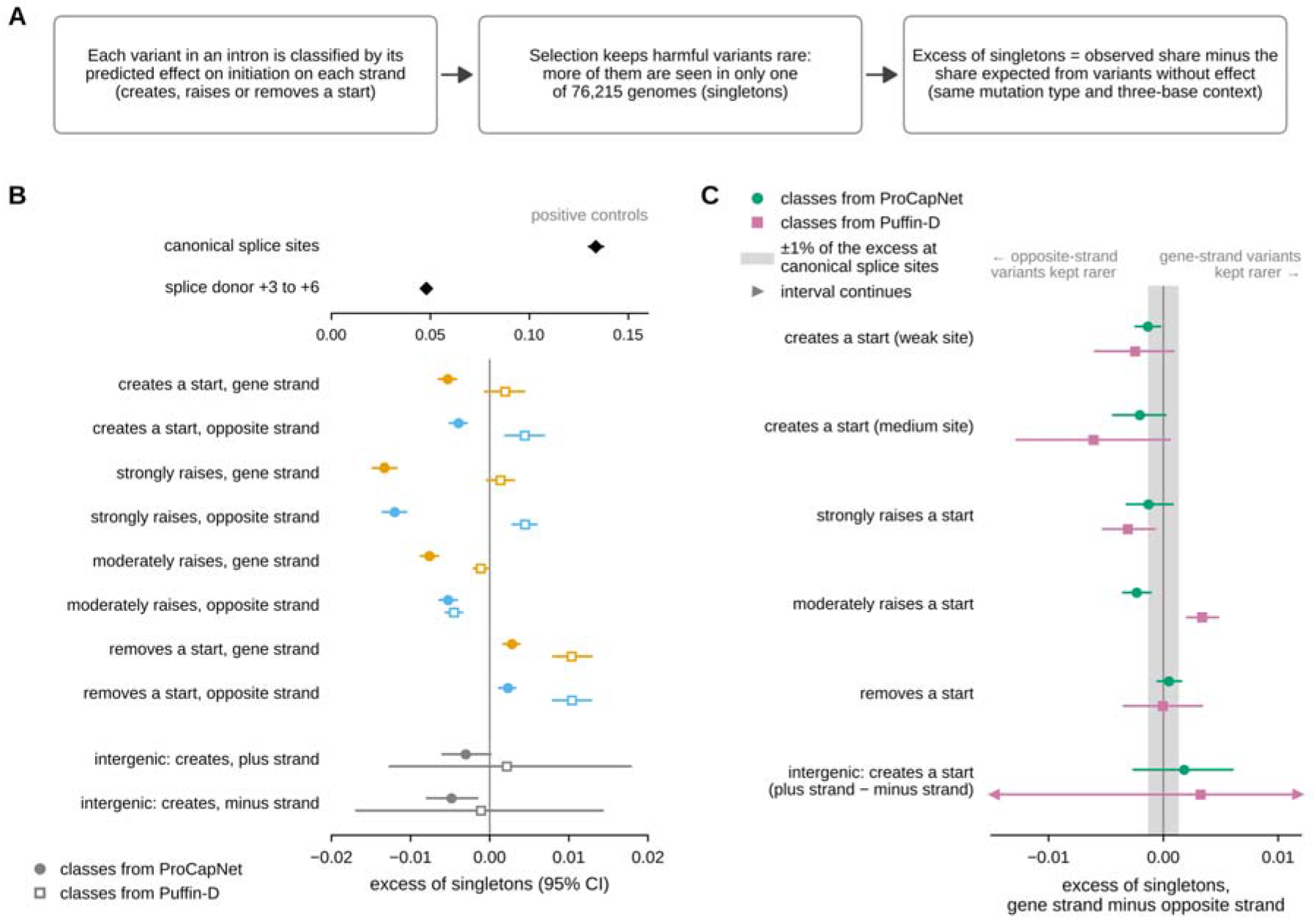
No sign of selection against new sense start sites. (A) Logic of the test: harmful variants are more often singletons, so a class under selection has an excess of singletons over variants with the same mutation type and three-base context but no predicted effect (oriented to the gene strand). (B) Excess of singletons for splice-site variants (positive controls, black diamonds, drawn on their own scale) and for intronic variants of protein-coding genes that create (weak sites), strongly or moderately raise, or remove (weak sites) a start on the gene strand (orange) or on the opposite strand (blue), and intergenic variants that create a weak site (plus and minus strand, gray). Filled circles, classes from ProCapNet; open squares, classes from Puffin-D; 95% CI from 1,000 bootstrap resamples of genes (intergenic: 1-Mb blocks). (C) Gene-strand minus opposite-strand excess of singletons for five types of effect, and plus-strand minus minus-strand excess for intergenic variants that create a start, with classes from ProCapNet (green circles) and Puffin-D (pink squares), with 95% CI; arrowheads, interval continues beyond the axis. For classes that create or raise a start, a positive difference would indicate selection against sense starts; gray band, ±1% of the excess of singletons at canonical splice sites, that is, one hundredth of the splice-site value in B (not one percentage point).

We first checked that the test detects selection where selection is known to act. Variants at canonical splice sites had a large excess of singletons, and variants at positions 3–6 of splice donor sites a smaller one (Fig. 4B). Classes of variants that gain or lose a start, in contrast, showed only small excesses or deficits of singletons, of similar size on both strands and also in intergenic DNA, where neither strand is special. These small values may reflect differences in mutation rate that the three-base context does not capture. Such differences should be similar on the two strands, so the informative comparison is between the strands: if new sense starts are harmful, variants that create a start on the gene strand (a sense start) should have a larger excess of singletons than variants that create a start on the opposite strand (an antisense start).

They did not. In the ProCapNet classes, variants that create a sense start had a slightly smaller excess of singletons than variants that create an antisense start (difference −0.0014, 95% CI −0.0025 to −0.0002; Fig. 4C). This small difference should not be over-interpreted: differences of this size and larger also occurred between the plus and minus strands of intergenic DNA, where no difference is expected (for ProCapNet variants that remove a medium site, −0.0099, −0.0173 to −0.0022; Supplemental Table S1). The difference was also zero or negative for ProCapNet variants that create medium sites or strongly raise a start, and for Puffin-D variants that create or strongly raise a start (Fig. 4C). Puffin-D predicts about ten times fewer created starts than ProCapNet, so the Puffin-D estimates for these classes rest on fewer variants and are less precise. These results held after excluding CpG sites and after matching for local GC content (Supplemental Fig. S7). In intergenic DNA, variants that create a start did not differ between the strands.

How strong a selection can these data rule out? To express this in familiar units, we compared the strand differences with the excess of singletons at canonical splice sites, where selection is strong. In the main analysis, for every class that creates or strongly raises a start, with either model, the upper end of the 95% CI lay below one hundredth of the splice-site excess. In other words, if variants predicted to create a sense start raised the excess of singletons by 1% or more of the excess at canonical splice sites, the test would have detected it. Without CpG sites (with or without matching for GC content), the upper end of the CI for Puffin-D variants that create a weak site reached about 1.5% of the splice-site excess.

Two further observations are described in the Supplemental Results. One class of variants that Puffin-D predicts to raise a start moderately had more singletons on the gene strand, but the corresponding ProCapNet classes showed the opposite; and the two models agreed poorly on which variants create sense starts, so the conclusion does not rest on one model (Supplemental Fig. S8).

Intronic variants of protein-coding genes in the two compared classes, all autosomes: ProCapNet, 2,610,417 (create a weak site), 911,755 (strongly raise a start) and 2,249,974 (moderately raise a start); Puffin-D, 275,674, 642,746 and 1,800,949. gnomAD v4.1 genomes, 215.5 million single-nucleotide variants with allele number at least 80% of the maximum.

## Discussion

We asked why, at elements inside human genes that start transcription in both directions, the start that points in the direction of the gene is weaker. Of the three explanations that we considered (Fig. 1C), the data favor the third: the bias is a property of the active elements (Fig. 5). The bias was present in both cell lines, and it remained when we removed pairs near annotated transcript starts or pairs that overlap transposable elements. It did not change detectably with the activity of the host gene. Sequence models that had never seen the chromosome of a pair predicted 30–42% of the bias from the local sequence of the element. In contrast, introns as a whole show little or no depletion of sequences able to start sense transcription, and variants that create such sequences are not detectably removed from the population.

**Figure 5.**
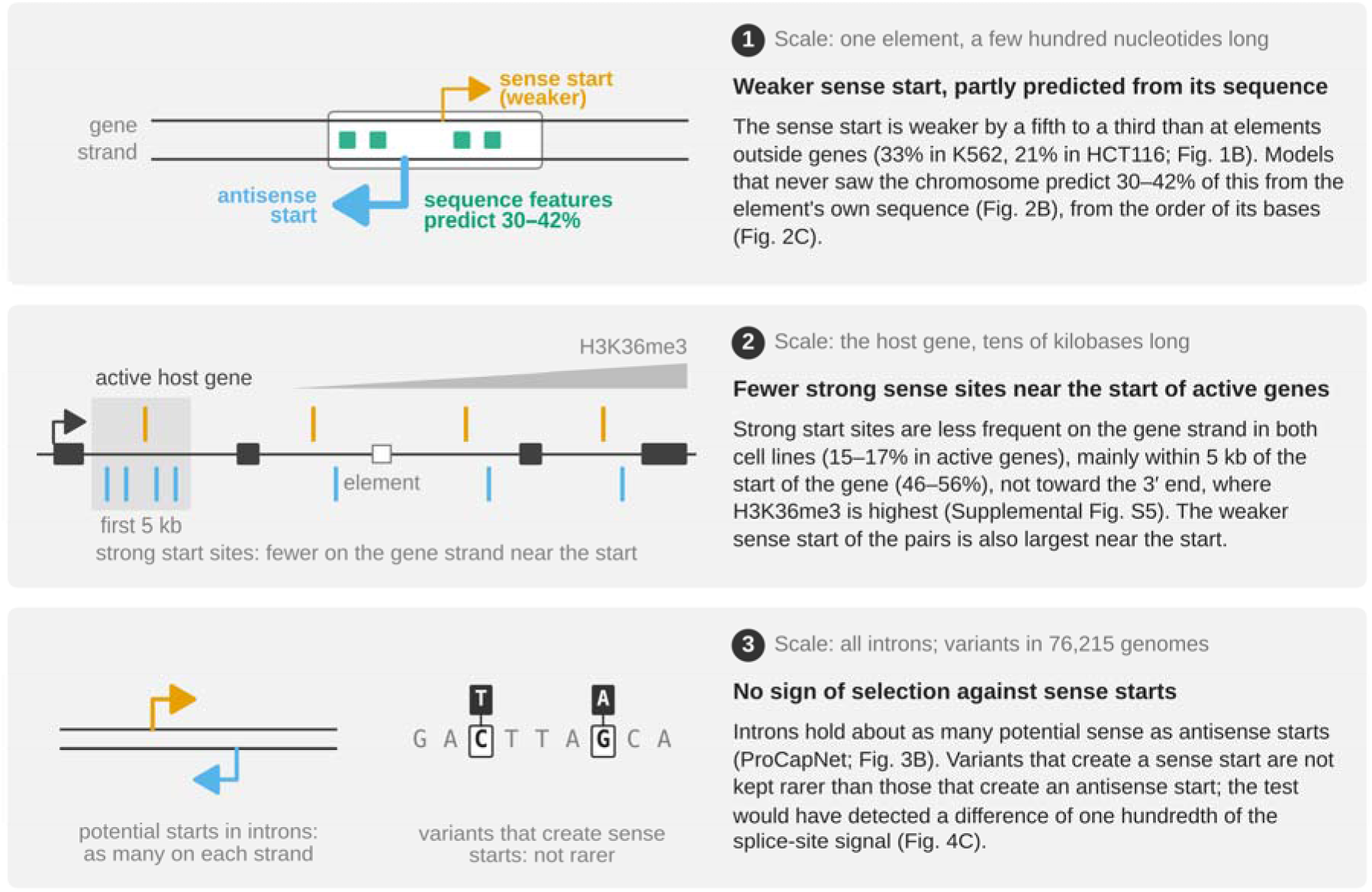
The conclusions at three scales. Schematic; no data are plotted. (1) A single element, which spans a few hundred nucleotides: the sense start (orange) is weaker than the antisense start (blue), by a fifth to a third relative to elements outside genes (Fig. 1B), and 30– 42% of this difference is predicted from the element’s own sequence (green boxes; Fig. 2B, C). (2) The host gene, which spans tens of kilobases (dark boxes, exons; white box, the element in 1): strong measured start sites (bars above the line, gene strand; below the line, opposite strand) are less frequent on the gene strand, mainly within the first 5 kb of active genes (shaded), whereas H3K36me3 (gray wedge) rises toward the 3′ end (Fig. 3F; Supplemental Fig. S5). (3) All introns and variants in 76,215 genomes: introns contain about as many potential sense as antisense starts (Fig. 3B), and variants that create a sense start (a base that differs between people, dark box above the reference base) are not kept rarer than variants that create an antisense start (Fig. 4C).

What in the elements could make the sense start weaker? Known directional elements, such as the TATA box and the YY1 motif (Dudnyk et al. 2024; Cochran et al. 2024), were not more common at the antisense start than at the sense start in a pre-specified test (Supplemental Fig. S2), so they do not explain the bias. In K562, our shuffling experiment shows where the information used by ProCapNet lies: in the order of the bases rather than in their composition, within a few hundred nucleotides that span both starts (Fig. 2C). This conclusion concerns only the part of the bias that the models predict (30–42%); the rest may come from sequence features that the models miss, from DNA outside the 2.1 kb that ProCapNet sees, or from factors other than the DNA sequence. Because models trained on colorectal cells predicted the bias in both cell lines, but models trained on K562 only in K562, the relevant features may differ between cell types, as does the activity of the elements themselves (ENCODE Project Consortium 2020). Which features these are remains to be found; one way to find them is to compare the contributions of individual bases to the ProCapNet prediction (Cochran et al. 2024) between the sense and antisense halves of intronic and intergenic elements. Finally, because PRO-cap captures RNA while it is still being made (Kwak et al. 2013), the bias arises when transcription starts or shortly afterwards. The bias is therefore distinct from the lower stability of antisense RNA, which is set by signals further downstream (Almada et al. 2013; Ntini et al. 2013).

Strong measured start sites point to a second influence on sense starts. In both cell lines and with a second assay, strong sites were less frequent on the gene strand, more so in the more active genes, and in K562 Puffin-D, which sees the host gene, partly reproduced this pattern, whereas ProCapNet, which sees only local sequence, did not. Mechanisms that suppress initiation inside transcribed genes are known: histone methylation by Set2 in yeast (Carrozza et al. 2005), DNA methylation recruited by H3K36me3 in mouse embryonic stem cells (Neri et al. 2017) and the H3K36 methyltransferase SETD2 in human cells (Carvalho et al. 2013). The deficit, however, does not appear to follow H3K36me3, the histone mark common to these mechanisms: this mark rises toward the 3′ end of active genes (Bannister et al. 2005), whereas in both cell lines, and with GRO-cap, the deficit of strong sense sites lay in the 5′ fifth of genes and was absent in the 3′ fifth. The link with host activity appeared for single strong sites but not for the start-site pairs. The two results need not conflict, because the pair analysis compares the two starts of one element, whereas the single-site analysis counts strong sites on each strand of all introns. Our test of DNA methylation gave no clear answer, and too few strong sites lay inside H3K36me3 peaks for a direct test. In GRO-cap data, a secondary analysis suggested a larger deficit inside peaks (22.2 percentage points, 95% CI −0.3 to 39.8; Supplemental Methods). Marks of elongation may therefore still contribute, but they are unlikely to explain the main part of the deficit, which lies near the start of genes.

The position of the deficit points instead to the region just downstream of gene promoters: the deficit of strong sense sites was confined to the first 5 kb of active genes, and the weaker sense start of the pairs was largest there. The excess of antisense over sense strong sites in this region may be related to downstream antisense transcription, which in a human cell line starts inside the gene within 2 kb of 28% of gene transcription start sites, often at the downstream edge of a CpG island, although an island is not required (Lavender et al. 2016). Because we compare the two strands, we cannot tell whether sense starts are suppressed near promoters or antisense starts are favored there. Unannotated promoters of the host gene would add strong sites mainly on the gene strand, so they would work against the deficit rather than create it.

The first explanation, selection against sequences able to start sense transcription, predicts that introns as a whole contain fewer potential sense starts. Puffin-D at first suggested exactly that, but this predicted deficit grew when the DNA of introns was shuffled and followed the transcription signal that PRO-cap records along active genes. Puffin-D sees 100 kb of DNA (Dudnyk et al. 2024), which can include the promoter of the host gene, and it appears to have learned where genes are transcribed. ProCapNet, which sees only 2.1 kb of local sequence, predicted little or no depletion of potential sense starts in real introns compared with shuffled introns of the same composition, and the strand differences that it predicted lay within the range of the intergenic control. Neither model therefore gives clear evidence that introns as a whole contain fewer potential sense starts. This conclusion concerns what intronic DNA could do, not which start sites the cells use; the deficit of strong measured sense sites described above is seen in the data but is not predicted from local sequence.

The population test looked for selection directly, and it found none: with either model, variants that create a sense start had an excess of singletons equal to, or slightly smaller than, that of variants that create an antisense start. The test readily detected selection at splice sites, and in the main analysis even a difference one hundredth the size of the splice-site excess, in the direction of selection, would have been detected. The two models classified largely different variants as creating a start, yet both gave the same result. The test has two limits. First, it detects only selection strong enough to change allele frequencies in recent human history; weak selection over long evolutionary times could escape it, but should have left fewer potential sense starts in introns across the genome, and ProCapNet found no such depletion. Second, each class of variants defined by the predictions of a model (for example, variants predicted to create a sense start) also contains variants that the model predicts wrongly, and these variants dilute any signal. Together, the census and the population test argue against a general removal of sense starts from human introns, although they do not exclude selection at particular genes or elements.

The second explanation, a mutational shift in base composition, is not supported either. The gene strand of introns does carry an excess of G and T over A and C (Green et al. 2003; Polak and Arndt 2008). However, a regression on this composition predicted none of the bias of the pairs, and for strong single sites the two models responded to this composition in opposite directions.

Two practical lessons follow. PRO-cap positions with few reads inside genes follow the transcribed strand of the host gene and of its neighbors, so comparisons of the two strands should use high read thresholds or genes without active neighbors. A sequence model that sees a long stretch of DNA can learn where genes are transcribed, so such comparisons also need a model with a short input, held-out chromosomes, an intergenic control and a check against measured data; even then, strand differences of a few percent should not be interpreted.

Our study has limitations. The start-site pairs come from two cell lines, and so do the strong measured start sites, but the comparison of measured with predicted sites (Fig. 3D) and the test of DNA methylation used K562 data only. The analyses of distance from the start of genes, of CpG islands and of activity near the start were chosen after seeing the data; of their findings, only the concentration within the first 5 kb was then confirmed in a pre-specified test (with K562 GRO-cap data), and the others need confirmation in other cell types. In K562, the pairs, the single sites and the training data of the K562 ProCapNet models all come from one PRO-cap experiment, so holding out chromosomes does not remove technical features shared within that experiment; the Caco-2 models, which share no data with either cell line, gave similar shares (41% in K562 and 30% in HCT116). The HCT116 PRO-cap data come from untreated cells of a line engineered for auxin-inducible degradation of CTCF, and we cannot exclude that the engineering affected initiation. Both cell lines come from cancers, whereas the population test concerns selection in the germline. Finally, the effects of variants are predictions, not measurements.

In summary, the weaker sense start at intronic start-site pairs appears to be a property of the elements that are active inside genes, partly predictable from their sequence and largest near the start of the gene. It does not appear to result from a genome-wide removal of sequences able to start sense transcription from introns (Fig. 5). Human introns seem to keep about as much sequence able to start sense transcription as antisense transcription, so whatever limits sense initiation seems to act at the active elements or in their surroundings. Where such control fails, this potential may become visible: in mouse embryonic stem cells, for example, loss of DNA methylation inside genes leads to spurious initiation within genes (Neri et al. 2017). Why active intronic elements should be built this way remains open. One possible reason is interference with the host gene: transcription from intragenic enhancers can lower the expression of their host genes (Cinghu et al. 2017). If such interference is stronger for transcription in the sense direction, which our data do not test, elements whose stronger start points against the gene might be better tolerated inside active genes. Endogenous retroviruses show a similar bias: in human introns they lie more often in the antisense orientation, probably because sense copies, with their splice and polyadenylation signals, are removed by selection (van de Lagemaat et al. 2006). Selection of this kind would act on relatively few elements and could escape the genome-wide tests used here. Testing this idea will require following the elements across cell types and species, and asking whether variants inside active elements that strengthen the sense start are kept rare.

## Methods

### Data

We analyzed public data only, on the GRCh38 genome and the GENCODE v47 annotation (Mudge et al. 2025), restricted to the 22 autosomes (the start-site pairs also include 42 pairs on chromosome X in K562 and 38 in HCT116; leaving them out changed the bias by about 1 percentage point). All ENCODE files (ENCODE Project Consortium 2020) came from the ENCODE portal (https://www.encodeproject.org; Luo et al. 2020). K562 PRO-cap came from experiment ENCSR261KBX, generated with the CoPRO protocol (Tome et al. 2018), and HCT116 PRO-cap from experiment ENCSR165QKR, done in untreated cells of an HCT116 line engineered for auxin-inducible degradation of CTCF. ProCapNet models came from the ENCODE annotation sets ENCSR740IPL (K562) and ENCSR182QNJ (Caco-2). For single start sites, the K562 signal was the 5′-end signal released with the K562 ProCapNet models, derived from the same experiment, and the HCT116 signal was the default ENCODE processing. K562 GRO-cap with TAP came from GEO GSM1480321 (Core et al. 2014), mapped to hg19 by the authors; we transferred the counts to hg38 (Supplemental Methods). H3K36me3 replicated peaks (K562 and HCT116) and K562 CpG methylation from whole-genome bisulfite sequencing also came from ENCODE. Variants were single-nucleotide variants passing quality filters in the 76,215 genomes of gnomAD v4.1 (https://gnomad.broadinstitute.org; Guez et al. 2026), which include the 76,156 genomes of gnomAD v3.1.2 (Chen et al. 2024). Transposable elements and CpG islands came from the RepeatMasker and cpgIslandExt tracks of the UCSC Genome Browser (https://hgdownload.soe.ucsc.edu). File identifiers, checksums and download dates are given in the Supplemental Methods.

### Start-site pairs and their asymmetry

Pairs of start sites were the bidirectional peaks called by PINTS (Yao et al. 2022) in K562 (ENCFF819CPA) and, with the same settings, in HCT116, and every called pair was used, whatever its confidence level; pairs were classified with the rules of our earlier analysis of intronic initiation (MW Szcześniak and E Wanowska, unpubl.). A pair was intronic when both starts lay in one intron of the reference transcript of a protein-coding gene (MANE Select, else Ensembl canonical, else the longest coding transcript), at least 1 kb from every exon of the gene’s protein-coding transcripts; pairs in readthrough genes or inside genes on both strands were left out, and of overlapping genes on one strand the shortest was taken as the host. A pair was intergenic when no protein-coding transcript lay within 5 kb of it. Other pairs were not used. In intronic pairs, the start on the strand of the host gene was the sense start; in intergenic pairs, the plus-strand start was called sense, an arbitrary choice that centers the reference on zero. S and A are the PRO-cap reads within 50 nt of the sense and the antisense start, and the asymmetry of a pair is DI = log2((S + 1)/(A + 1)). A host gene was active when its annotated start carried at least five PRO-cap reads.

For the matched comparison, intronic pairs were taken in random order, and each was matched, without replacement, to the intergenic pair of nearest read depth (S + A) among those not yet used, provided that the depths differed by no more than 10%. The difference d between the median DI of intronic pairs and that of their matched intergenic pairs was converted to the percentage by which the sense start is weaker, 100 × (1 − 2). Confidence intervals came from 10,000 resamples of host genes, each with its matched intergenic pairs, and matched pairs were compared with a two-sided Wilcoxon signed-rank test. The depth bands, the analyses by host activity and H3K36me3, the removal of pairs near annotated transcript starts and the analyses of transposable elements are described in the Supplemental Methods.

### Sequence models and predictions for pairs

Puffin-D (Dudnyk et al. 2024; https://github.com/jzhoulab/puffin, commit 9bc81eb) was run on 100-kb windows centered on each pair. Puffin-D was trained on all chromosomes except 8 and 9 (test) and 10 (validation). ProCapNet (Cochran et al. 2024) was run with the seven K562 models released by ENCODE (2,114-nt input), each trained without one group of test chromosomes (Supplemental Fig. S1). Each pair was scored with the ProCapNet model that had not seen its chromosome (held-out prediction) and, for comparison, with the other six models; the same was done with the seven Caco-2 models, which use the same chromosome groups. Each window was scored as given and as its reverse complement, and the two predictions were averaged. The predicted asymmetry of a pair was computed from the predicted signal within 50 nt of each start (Supplemental Methods).

Predictions are compressed relative to measurements, so they were calibrated before comparison: measured DI was regressed on predicted DI by least squares through the origin, among intergenic pairs taken once in each orientation. The share of the bias predicted from sequence was the calibrated mean predicted DI of intronic pairs in active hosts divided by their mean measured DI. Confidence intervals came from 2,000 resamples of host genes for intronic pairs and, independently, of 1-Mb genomic blocks for intergenic pairs. For the shuffles, the 2,114 nt seen by ProCapNet were shuffled with an algorithm that keeps the count of every base and every dinucleotide (Altschul and Erickson 1985), in the whole window, in the element (from 100 nt before the first start to 100 nt after the second) or in the two flanks, five times per pair. Base composition was modeled by a linear regression of DI on the strand skews of T over A and of G over C within 300 nt of the pair, fitted on intergenic pairs (Supplemental Methods).

### Genome-wide census of potential start sites

Every position of the 22 autosomes was scored on both strands with both models, averaging the two orientations; ProCapNet used, for each chromosome, the model that had not seen it, and Puffin-D used 100-kb windows every 50 kb, keeping the central 50 kb of each. Eligible intronic positions lay in protein-coding or lncRNA genes, excluding exons and 100 nt on each side, 1 kb around every annotated transcript start, positions covered by another gene and positions with an unknown base. Intergenic control positions lay more than 10 kb from any annotated gene and were grouped into 1-Mb blocks, with the plus strand standing for the gene strand.

Thresholds for weak, medium and strong sites were the 0.999, 0.9997 and 0.9999 quantiles of all predictions for chromosomes 8 and 9, and positions at or above a threshold within 50 nt of each other were merged into one site. The strand ratio of a set of genes was the sum of gene- strand sites divided by the sum of opposite-strand sites, and the deficit was 100 × (1 − ratio); confidence intervals came from 2,000 resamples of genes or blocks. For the shuffle test, eligible positions were shuffled with the same dinucleotide-preserving algorithm in pieces of up to 500 nt, separately within each gene and repeat status, and scored again.

### Measured start sites in K562 and HCT116

Measured start sites were positions with at least 1, 3 or 10 PRO-cap reads (18.7 million reads in total) at the eligible positions, merged within 50 nt; sites with at least 10 reads are called strong. The activity of a gene was the number of reads on its strand within 250 nt of any of its annotated transcript starts: silent, 0; weakly active, 1–4; active, 5 or more, divided into thirds of genes. This definition differs from the one used for host genes of pairs, which counts reads at one annotated start. In HCT116, strong sites had at least 24 reads, the K562 threshold scaled to the read depth. A site was inside an H3K36me3 peak when the position of the site with the most reads lay in a replicated ENCODE peak, and each gene was divided into fifths along the annotated gene. For K562 GRO-cap, strong sites had at least 14 reads, the threshold scaled in the same way. The analyses of silent genes, of DNA methylation, of distance from the start of genes and of CpG islands are described in the Supplemental Methods.

### Population test

We used the single-nucleotide variants of gnomAD v4.1 genomes at sites where the number of genotyped alleles was at least 80% of the maximum (152,430): all protein-coding intronic variants and random subsets of lncRNA intronic (30%) and intergenic (20%) variants at eligible positions, 215.5 million variants in total. Each variant was scored with the reference and with the alternative allele, reading the prediction within 50 nt on each strand, with Puffin-D and, separately, with the held-out ProCapNet model. A variant created a start when, on one strand, the reference prediction was below the weak or medium threshold and the alternative prediction was at or above it, and it removed a start in the reverse case. Strong and moderate raises were defined by cut-offs on the change on one strand with little change on the other strand, and neutral variants changed the prediction little on both strands (cut-offs in the Supplemental Methods).

The expected share of singletons in a class was the mean, over its variants, of the share of singletons among neutral variants with the same three-base context and alternative base, oriented to the gene strand (192 contexts), and the excess of singletons was the observed minus the expected share (Lek et al. 2016). Positive controls were variants at canonical splice sites and at positions 3–6 of splice donor sites. Differences between classes were bootstrapped over genes (1-Mb blocks for intergenic DNA; 1,000 resamples). The analysis was repeated without CpG sites, with contexts split by local GC content, and with both.

### Statistical analyses

Confidence intervals are 95% percentile bootstrap intervals; the resampled units and the numbers of resamples are given in the text and figure legends, and random seeds were fixed. Tests are two-sided, and P values are not corrected for multiple testing. Conclusions rest on the confidence intervals of four primary comparisons: the depth-matched asymmetry, the share predicted by held-out ProCapNet models, the gene-strand deficit of potential and measured start sites, and the sense-minus-antisense difference in the excess of singletons. The follow-up of strong measured sites (HCT116, H3K36me3 peaks and fifths of genes) was specified before the data were seen, as was its repetition with K562 GRO-cap data (the deficit, the 3′ minus the 5′ fifth, and the first 5 kb minus beyond); the analyses of distance from the start of genes, of CpG islands and of activity near the start were chosen afterwards and are labeled exploratory. The other analyses are consistency checks.

### Software availability

All analyses were run with scripts written for this study in Python 3.11 (NumPy, pandas, SciPy, pyfaidx, pyBigWig and PyTorch), and figures were drawn with matplotlib directly from the result tables. The scripts and the software environment are available at https://github.com/michal-szczesniak/intronic-start-asymmetry, are archived together with the result tables at Zenodo (https://doi.org/10.5281/zenodo.24161109) and are provided as Supplemental Code; versions are listed in the Supplemental Methods.

## Data access

This study generated no new data. All data analyzed are public, and their accessions are listed in the Methods and the Supplemental Methods: ENCODE (ENCSR261KBX, ENCSR165QKR, ENCSR000AKR, ENCSR091QXP, ENCSR740IPL, ENCSR182QNJ, ENCSR765JPC), GEO (GSE219647, GSE219542), gnomAD v4.1, GENCODE v47 and the UCSC RepeatMasker track. Supplemental Table S1 gives the results of all robustness analyses, of the DNA methylation analysis and of the population test for every class of variants, with the result file of each row; Supplemental Data also list every start-site pair with its read counts, its host gene, the predictions of each model and the other per-pair measures used in this study. Analysis scripts and result tables are archived at Zenodo (https://doi.org/10.5281/zenodo.24161109); the scripts are also available at https://github.com/michal-szczesniak/intronic-start-asymmetry.

## Acknowledgments

Computations were performed at the Poznań Supercomputing and Networking Center (PCSS), computational grant pl0074-02. Claude (Anthropic) was used to help write and debug analysis code, cross-check numbers in the text against the analysis output, and edit the text. The authors ran and verified all code, checked all text, numbers and citations, and take full responsibility for the content of the article.

## Author contributions

Michał W. Szcześniak: Conceptualization, Methodology, Software, Formal analysis, Investigation, Data curation, Visualization, Writing – original draft, Writing – review & editing, Supervision, Project administration. Elżbieta Wanowska: Conceptualization, Writing – original draft, Writing – review & editing, Funding acquisition.

