## Supplemental Material for "No evidence of genome-wide selection against sense transcription starts in human introns"

### Contents

- Supplemental Results
- Supplemental Methods
- Supplemental Figures
  - Supplemental Figure S1. Which model saw which chromosome, and the share of the asymmetry predicted under three calibrations
  - Supplemental Figure S2. Known core promoter elements do not distinguish the two starts of intronic pairs
  - Supplemental Figure S3. In silent genes, the strand difference of weak measured sites follows the strand reached by transcription from active neighbors
  - Supplemental Figure S4. Gene-body DNA methylation and strong measured start sites in K562
  - Supplemental Figure S5. The gene-strand deficit of strong measured sites lies near the start of genes
  - Supplemental Figure S6. Strand ratios of strong and weak start sites by the keto skew of the gene strand
  - Supplemental Figure S7. The strand comparison of the population test under four definitions of the expectation
  - Supplemental Figure S8. ProCapNet and Puffin-D classify largely different variants
- Supplemental References

Other supplemental files, provided separately: Supplemental Table S1 (results of all robustness analyses, of the DNA methylation analysis and of the population test for every class of variants, with the result file of each row); Supplemental Data (every start-site pair with its read counts, its host gene, the predictions of each model and the other per-pair measures); Supplemental Code (the analysis scripts and the software environment, also archived at Zenodo, https://doi.org/10.5281/zenodo.24161109).

### Supplemental Results

These analyses support the main Results.

#### Start-site pairs are rarely alternative promoters

Could these pairs simply be alternative promoters of their host genes? This is unlikely. Among all intronic pairs, the start closer to an annotated transcript start lay a median of 4–5 kb from it, and only 6–8% of the pairs had a start within 100 nt of one (5.3–6.5% of the pairs in active hosts; Supplemental Table S1). Intergenic pairs, in contrast, often sit at the annotated start of a non-coding transcript (37% in K562 and 28% in HCT116), so the reference group may be enriched for promoters. However, removing every pair, intronic or intergenic, with a start within 100 nt of an annotated transcript start did not change the result: the sense start was then weaker by 35% in K562 and 22% in HCT116, compared with 33% and 21% for all pairs. Removing pairs within 1 kb gave 32% and 19% (Supplemental Table S1).

#### Base composition, core promoter elements and transposable elements

Could base composition alone explain the bias? Within 300 nt of the pairs, the gene strand of introns carries more T than A (about 11% more) and more G than C (about 5–6% more), whereas intergenic pairs show no such difference. If composition caused the bias, these strand differences should predict the asymmetry of a pair. We fitted a linear regression that predicts the asymmetry of a pair from these two strand differences, using intergenic pairs, and tested it on pairs left out of the fit (cross-validation; Supplemental Methods). The regression explained less than 1% of the variation in asymmetry among intergenic pairs (cross-validated $R^2$ 0.002 in K562 and 0.006 in HCT116). Applied to intronic pairs, it predicted a slightly stronger, not a weaker, sense start (share −24% in K562 and −11% in HCT116; Fig. 2B). Adding base composition to the Puffin-D prediction in one linear model did not raise the share; the share fell slightly, from 35% to 29% in K562 (Supplemental Table S1). Base composition therefore does not explain the bias.

Within the element, the two starts could differ in known core promoter elements. To test this, we compared the sequence around the two starts of each depth-matched pair (1,610 pairs in K562 and 2,489 in HCT116; Supplemental Fig. S2A). As expected for precisely mapped starts, 75–85% of the starts lay at a pyrimidine followed by a purine, compared with about 20% at positions three or more nucleotides away. Before the analysis, we chose five elements: the TATA box, the initiator, the downstream promoter element and the YY1 motif in each orientation. For each element, we counted the starts that carried it at its usual position and subtracted the number expected from the base composition around the start. We then asked whether the sense and the antisense start of intronic pairs differed more than the two starts of their intergenic partners. None of the five elements did, in either cell line (two-sided bootstrap test over host genes; Holm-corrected $P \geq 0.19$ in K562 and $P = 1$ in HCT116; Supplemental Fig. S2B). Among the secondary measures, a TATA box with one mismatch was less common at the sense start in K562 (by 2.4 percentage points of starts, 95% CI 0.3–4.5; $P = 0.03$ without correction) but not in HCT116 (by 0.3 points, −1.0 to 1.7), and, relative to the intergenic partners, the DNA around the sense start was slightly poorer in G and C in both cell lines (by 0.7 and 0.5 percentage points; P = 0.03 and 0.007). Known core promoter elements therefore do not explain the bias, and the features that ProCapNet uses remain to be identified.

Transposable elements could create the bias indirectly. The start that points in the same direction as a transposable element might be stronger, and transposable elements in introns might mostly point against the gene. A third to two fifths of the pairs (34–40%), in introns and between genes alike, overlap a transposable element. In introns, these transposable elements point against the gene in about three cases out of five (63% in K562 and 60% in HCT116), slightly more often than all transposable elements in the bodies of the same genes (57%). Of the main classes, LTR retrotransposons had the clearest effect on direction: the stronger start pointed in the same direction as the transposable element (Supplemental Table S1). However, removing every pair whose element overlaps a transposable element left the bias almost unchanged (31% in K562, 95% CI 11–47%; 25% in HCT116, 14–35%). Removing every pair that overlaps any repeat gave similar estimates (25% and 22%), although with almost half of the pairs removed, the K562 interval then included zero (−1 to 41%). Transposable elements therefore do not explain the bias.

### Weak measured sites in silent genes

The transcription signal probably also explains a puzzling result in silent genes, which have no reads at their own starts. In these genes, measured sites with at least one PRO-cap read were 28% less frequent on the gene strand than on the opposite strand. This deficit depended on transcription that enters the silent gene from an active neighboring gene (Supplemental Fig. S3). When the transcription of the neighbor ran along the opposite strand of the silent gene, the gene strand showed a large deficit; when it ran along the gene strand, the gene strand showed an excess; and silent genes without active neighbors showed no deficit. Sites with few reads therefore appear mostly to mark where transcription passes, not where it starts.

### DNA methylation and strong measured sites

We asked whether DNA methylation inside genes, which can prevent spurious initiation (Neri et al. 2017), is linked to the deficit of strong sense sites. More active genes are also more methylated (Supplemental Fig. S4A), so we compared genes with high and low methylation only within groups of genes with similar activity (Supplemental Methods). The planned comparison, which pools the strong sites of all activity groups, showed no difference in the deficit (4.5 percentage points, 95% CI −12.9 to 20.7). The two methylation groups, however, draw their strong sites from different genes. In the highly methylated group, most strong sites lie in the least active genes, which show no deficit; in the weakly methylated group, few do (Supplemental Fig. S4C). The pooled comparison is therefore dominated, in the highly methylated group, by genes without a deficit. A comparison that gives each activity group the same weight, which we chose after seeing the data, suggested a moderately larger deficit in more methylated genes (24.5 percentage points, 95% CI 1.4 to 48.0), but this estimate is uncertain (Supplemental Fig. S4D). We therefore found no clear link between DNA methylation inside genes and the deficit of strong sense sites.

### Strong sites near the start of genes: CpG islands and gene activity

Near the start of a gene, most strong sites and pairs lay within 2 kb of a CpG island. To separate the two factors, we pooled both cell lines (Supplemental Methods). More than 2 kb from any CpG island, the strand difference was still larger in the first 5 kb than farther away (strong sites, 26% compared with 4%, difference 22.5 percentage points, 95% CI 3.8–38.9, $P = 0.016$; pairs, 45% compared with 14%, difference 30.6, 1.7–54.1, $P = 0.04$), whereas CpG islands farther from the start did not detectably raise the strand difference (differences 3.3, −11.0 to 16.3, and 14.0, −15.3 to 33.5; Supplemental Fig. S5D). For strong sites, the excess in the first 5 kb was similar in all three thirds of gene activity (most minus least active third, 8.0 percentage points, −15.4 to 32.2); for the pairs, this comparison was too uncertain to interpret (17.5, −22.3 to 63.4; Supplemental Fig. S5E). These analyses were chosen after seeing the data and are exploratory.

### A Puffin-D class of variants and the agreement between the two models

One class departed from the overall pattern of the population test. Variants that Puffin-D predicts to raise a start moderately had a larger excess of singletons on the gene strand than on the opposite strand (difference 0.0034, 95% CI 0.0020 to 0.0049; Fig. 4C), also without CpG sites and with matching for GC content (Supplemental Fig. S7). The corresponding ProCapNet classes, however, showed the opposite (−0.0023, −0.0036 to −0.0010). Moreover, this Puffin-D class is the only class that is much larger on the gene strand than on the opposite strand (1.07 versus 0.73 million variants), and the Puffin-D predictions on the gene strand include the transcription signal (Fig. 3D). A moderate gain predicted by Puffin-D therefore need not mark a new start, and what this excess reflects is unclear.

The two models agree poorly on which variants matter. On chromosomes 8 and 9, ProCapNet classified about 123,000 intronic variants as creating a sense start and Puffin-D about 12,000, with only 726 in common (Supplemental Fig. S8). The two models thus reached the same conclusion from largely different variants, so the conclusion does not depend on one particular model. Both models, however, make errors: each class also contains variants that do not change initiation in reality, and these variants dilute any signal of selection. The limits given in the main text therefore apply to the variants as the models classify them; selection on the variants that truly create a sense start could be stronger than these limits suggest. Results for lncRNA introns, based on a 30% sample of variants, were imprecise (Supplemental Table S1).

### Supplemental Methods

#### Data sources

No cell lines or organisms were handled. Data came from K562 (chronic myelogenous leukemia) and HCT116 (colorectal carcinoma) cells profiled by ENCODE (ENCODE Project Consortium 2020), and from ProCapNet models trained on K562 and Caco-2 (colorectal adenocarcinoma) PRO-cap, and from Puffin-D (below). K562 PRO-cap came from experiment ENCSR261KBX, generated with the CoPRO protocol (Tome et al. 2018; GEO GSE219647). HCT116 PRO-cap came from experiment ENCSR165QKR (GEO GSE219542), done in untreated cells of an HCT116 line engineered for auxin-inducible degradation of CTCF (CTCF tagged with a degron, and the rice auxin receptor OsTIR1, LOC4335696, introduced). H3K36me3 peaks were the replicated ENCODE peak files ENCFF520LHY (experiment ENCSR000AKR, K562) and ENCFF029GQD (ENCSR091QXP, HCT116), identified by their md5 checksums. ProCapNet models came from ENCODE annotation sets ENCSR740IPL (K562) and ENCSR182QNJ (Caco-2; models ENCFF575XTL, chromosome splits ENCFF603POZ). The measured K562 signal used for single start sites was the observed 5′-end signal released with the K562 ProCapNet models (ENCFF798GNW, plus strand; ENCFF662SHP, minus strand), derived from experiment ENCSR261KBX. In K562, therefore, the pairs, the single start sites and the training data of the K562 ProCapNet models all come from one PRO-cap experiment. CpG methylation of K562 came from ENCODE whole-genome bisulfite sequencing (ENCSR765JPC): files ENCFF660IHA and ENCFF328NMN of the experiment's default analysis and, for comparison, files ENCFF867JRG and ENCFF721JMB of an earlier processing of the same reads with an older ENCODE pipeline, identified by their md5 checksums. K562 GRO-cap came from GEO (GSM1480321; see Measured start sites below).

#### Genome, annotation and databases

All analyses used the GRCh38 (hg38) genome and the GENCODE v47 annotation (Mudge et al. 2025), restricted to the 22 autosomes, except the start-site pairs, which include 42 depth-matched pairs on chromosome X in K562 and 38 in HCT116 (without them, the bias was 32.5%, 95% CI 11.6–45.3%, and 20.0%, 11.1–29.0%). ENCODE files were obtained from the ENCODE portal (https://www.encodeproject.org; Luo et al. 2020). Genetic variants were single-nucleotide variants passing quality filters in the genomes of gnomAD v4.1 (https://gnomad.broadinstitute.org; released 19 April 2024; 76,215 individuals, maximum allele number 152,430; Guez et al. 2026), which include the 76,156 genomes of gnomAD v3.1.2 (Chen et al. 2024). Transposable elements were taken from the RepeatMasker track of the UCSC Genome Browser (https://hgdownload.soe.ucsc.edu, table rmsk, downloaded on 23 October 2026), and CpG islands from the cpgIslandExt track of the same browser (downloaded on 10 November 2026, file dated 19 October 2022, md5 17da03fd8be6e33d2c09228df677a0f8; 26,866 islands on the 22 autosomes). Puffin-D was run from the authors' repository (https://github.com/jzhoulab/puffin, commit 9bc81eb of 3 March 2025).

### Start-site pairs and their asymmetry

Pairs of start sites were the bidirectional peaks called by PINTS (Yao et al. 2022) in K562 (ENCFF819CPA) and, with the same settings, in HCT116, and were selected as in our earlier analysis of intronic initiation (MW Szcześniak and E Wanowska, unpubl.). Each pair has one start on each strand, and the element is the region spanning the two starts (analyses that add a margin around it say so). Every called pair was used, whatever its confidence level. A pair was intronic when both starts lay in one intron of the reference transcript of a protein-coding gene (MANE Select, else Ensembl canonical, else the longest coding transcript), at least 1 kb from every exon of the gene's protein-coding transcripts; pairs in readthrough genes or inside genes on both strands were left out, and of overlapping genes on one strand the shortest was taken as the host. A pair was intergenic when no protein-coding transcript lay within 5 kb of it. Other pairs were not used. For intronic pairs, the start on the host gene's strand was the sense start. For intergenic pairs, the plus-strand start was called sense; this choice is arbitrary, so the reference is expected to be centered on zero. S and A are the PRO-cap reads within 50 nt of the sense and antisense start, and the asymmetry of a pair is the directionality index $DI = \log2((S + 1)/(A + 1))$. The host gene was called active when its annotated start carried at least five PRO-cap reads, as in the earlier analysis; the analysis of single start sites (below) measures the activity of a gene anew, from reads near any of its annotated starts, so the two definitions are not identical.

Read depth (S + A) was divided into five bands at the quintiles of the pooled intronic and intergenic pairs, and within each band the median DI of intronic pairs in active hosts was compared with that of intergenic pairs. For the matched comparison, intronic pairs were taken in random order, and each was matched, without replacement, to the intergenic pair of nearest depth among those not yet used, provided that the depths differed by no more than 10%. The difference d between the median DI of intronic pairs and that of their matched intergenic pairs was converted to the percentage by which the sense start is weaker, $100 \times (1 - 2^{d})$. Pairs in the same host gene are not independent, so confidence intervals were obtained by resampling whole host genes together with their matched intergenic pairs (10,000 resamples); in the depth bands, intergenic pairs were resampled in 1-Mb genomic blocks. Matched pairs were compared with a two-sided Wilcoxon signed-rank test. For comparison, confidence intervals were also computed by resampling single pairs, with groups compared by a two-sided Mann–Whitney U test (Supplemental Table S1). To check that the result does not depend on pairs at annotated promoters, the analysis was repeated, unchanged, after removing from both groups every pair with either start within 100 nt, or within 1 kb, of the start of any GENCODE transcript; with no pairs removed, this run reproduced the original tables exactly.

For the analysis of host activity, every intronic pair, in active or silent hosts, was matched to an intergenic pair of the same depth as above. The difference in DI between each intronic pair and its matched intergenic pair was summarized in groups of host activity: silent (no reads at the host's start), weakly active (1–4 reads) and three thirds of host activity among pairs in active hosts, so that each third holds the same number of pairs. The trend was measured by the Spearman correlation between host activity and this difference, with a 95% confidence interval from 10,000 resamples of host genes; the same resampling gave the confidence interval of the difference between active and silent hosts. A pair counted as marked by H3K36me3 when a peak of this mark lay within 1 kb of its midpoint.

### Sequence models and predictions for pairs

Puffin-D (Dudnyk et al. 2024) was run on 100-kb windows centered on each pair, and predicted PRO-cap signal was read on the plus and minus strand (output tracks 4 and 5). Each window was scored as given and as its reverse complement, the second output was mapped back to genomic coordinates, and the two were averaged, so that predictions do not depend on which strand the model sees. For each start, we took the highest prediction within 50 nt on its own strand, and the predicted asymmetry was the difference of the two values divided by ln 2. Puffin-D was trained on all chromosomes except 8 and 9 (test) and 10 (validation), which hold 16% of the matched K562 pairs and 14% of the matched HCT116 pairs; its predictions for most pairs therefore come from chromosomes it was trained on.

ProCapNet (Cochran et al. 2024) was run with the seven K562 models released by ENCODE (2,114-nt input, 1,000-nt output). Each model was trained without one group of test chromosomes (Supplemental Fig. S1). Each pair was scored with the model whose test group contained the pair's chromosome (held-out prediction) and, for comparison, with the six models for which this chromosome was not a test chromosome (train-fold prediction; predictions averaged). Both orientations were averaged as above, and the predicted asymmetry was log2 of the ratio of predicted counts within 50 nt of each start, with a pseudocount of 1, as for measured DI. The same was done with the seven ProCapNet models trained on Caco-2 PRO-cap, whose chromosome groups are identical to those of the K562 models.

Predictions are compressed relative to measurements, so they were calibrated before comparison. Among intergenic pairs, taken once in each orientation so that the reference is centered on zero, measured DI was regressed on predicted DI by least squares through the origin. The share of the asymmetry predicted from sequence was the calibrated mean predicted DI of intronic pairs in active hosts divided by their mean measured DI. As a check that needs no calibration, a rank-based share compared how far intronic pairs stand out from intergenic pairs in predicted and in measured DI (area under the curve minus 0.5, predicted divided by measured); this share is on a different scale. Calibration was also repeated within three bands of read depth (thirds of S + A among intergenic pairs), each intronic pair calibrated within its own band. Confidence intervals came from 2,000 resamples of host genes for intronic pairs and, independently, of 1-Mb genomic blocks for intergenic pairs.

For the pair-level shuffles, the 2,114 nt seen by ProCapNet around each pair were shuffled with the Altschul–Erickson algorithm (Altschul and Erickson 1985), which keeps the count of every base and every dinucleotide (runs of N left in place), in one of three regions: the whole window; the element, from 100 nt before the first start to 100 nt after the second; or the two flanks outside the element, each shuffled separately. Each region was shuffled five times per pair with different seeds, the predicted signal of the held-out K562 model was averaged over the five shuffles, and the share was calculated as above. The unshuffled sequence, scored in the same run, reproduced the earlier predictions to within 0.003 log2 units and gave the same shares.

#### Base composition

For each pair we computed, on the gene strand (for intergenic pairs, the plus strand) within 100, 300 and 1,000 nt of the pair, the skews (T − A)/(T + A) and (G − C)/(G + C) and the skews of six non-palindromic dinucleotide pairs. Linear models predicting DI from these features were fitted on intergenic pairs in both orientations and evaluated by five-fold cross-validation; the share was computed as for the sequence models, with 95% confidence intervals from 1,000 resamples of host genes and of intergenic pairs. Figure 2B shows the model with the two mononucleotide skews within 300 nt; all models are in Supplemental Table S1. To ask whether composition adds to a sequence model, the same features were added to the calibrated Puffin-D prediction in one linear model, fitted and evaluated in the same way.

#### Transposable elements

Transposable elements were RepeatMasker elements of the classes SINE, LINE, LTR, DNA, Retroposon and RC (a trailing question mark in the class was ignored). A pair overlapped a transposable element when its element, extended by 50 nt beyond each start, overlapped one; the transposable element with the largest overlap defined the class and the orientation, the latter relative to the host gene's strand for intronic pairs. As background, the orientation was counted for all transposable elements whose midpoint lies in the bodies (GENCODE gene spans) of the same host genes. The direction imposed by a transposable element was measured as log2((reads at the start on the element's strand + 1)/(reads at the other start + 1)). The depth-matched comparison was repeated, unchanged, without the pairs that overlap a transposable element, and without the pairs that overlap any RepeatMasker element.

#### Core promoter elements at the two starts

Each start of the depth-matched pairs (1,610 in K562 and 2,489 in HCT116) was read on its own strand from 60 nt before to 60 nt after the start, position 0 being the first transcribed base; for intergenic partners, the plus-strand start took the place of the sense start. Five elements were chosen before the analysis and counted as exact matches (IUPAC codes) at fixed positions: the TATA box, TATAWAWR starting between positions −36 and −24; the initiator, BBCABW with its A between −3 and +3; the downstream promoter element, RGWYV starting between +24 and +32; and the YY1 motif, AANATGGC on the transcribed strand (forward) or GCCATNTT (reverse), starting anywhere between −60 and +52. The expected share of starts carrying an element was its mean over 20 dinucleotide-preserving shuffles (Altschul and Erickson 1985) of the 121-nt window of the same start. For each element, the difference between the sense and the antisense start of intronic pairs (observed minus expected share) was compared with the same difference between the plus and the minus start of their intergenic partners. Confidence intervals and two-sided P values came from 2,000 bootstrap resamples of host genes, each intronic pair resampled together with its intergenic partner, and P values were Holm-corrected over the five elements within each cell line. Secondary measures were the TATA box with at most one mismatch, the G + C content and the number of CpGs between −50 and +50. As a check of the start positions, the share of starts at a pyrimidine–purine dinucleotide (positions −1 and 0) was computed at the given position and at positions shifted by up to 10 nt. The logos (Supplemental Fig. S2A) show the information content, $2 + \Sigma\, p \log_2 p$, without correction for sample size.

### Genome-wide maps of potential start sites

Every position of the 22 autosomes was scored on both strands. Puffin-D used 100-kb windows every 50 kb, keeping the central part of each, and ProCapNet used windows whose 1,000-nt outputs tile the genome, each chromosome scored by the model whose test group contained it; both were averaged over the two orientations. Eligible intronic positions were those inside protein-coding or lncRNA (long non-coding RNA) genes, excluding exons and 100 nt on each side, 1 kb around every annotated transcript start, positions covered by another gene on the opposite strand or by a second protein-coding or lncRNA gene on the same strand, and positions with an unknown base. Of the 14,267 protein-coding genes with eligible positions, Puffin-D tables lacked one gene (14,266 analyzed). Intergenic control positions lay more than 10 kb from any annotated gene and were grouped into 1-Mb blocks; in these blocks the plus strand stood for the gene strand, so their strand ratio is expected to be 1.

Thresholds for weak, medium and strong sites were the quantiles 0.999, 0.9997 and 0.9999 of all predictions for the real sequence of chromosomes 8 and 9, both strands (Puffin-D: 0.0439, 0.1432 and 0.3293; ProCapNet: 1.131, 2.658 and 5.848 predicted reads). Positions at or above a threshold were merged into one site when they lay within 50 nt of each other. For each gene we counted sites on the gene strand and on the opposite strand, and the strand ratio was the sum over genes of gene-strand sites divided by the sum of opposite-strand sites; deficits are reported as $100 \times (1 - \text{ratio})$. Counts were also made on non-repeat positions only (not soft-masked in the hg38 sequence). The summed signal ratio was the sum of predictions over all eligible positions of a set of genes on the gene strand divided by the same sum on the opposite strand. Confidence intervals came from 2,000 bootstrap resamples of genes or blocks.

For the shuffle test, eligible positions were divided into runs belonging to one gene, or to intergenic DNA, with one repeat status (soft-masked or not), and each run was shuffled in pieces of up to 500 nt with a random Eulerian-path algorithm that keeps the counts of all bases and dinucleotides (Altschul and Erickson 1985); the counts were checked to be identical before and after shuffling. All other positions were left unchanged, and the shuffled chromosomes were scored again with the same models and thresholds. Keto skew, $(G + T - A - C)/(A + C + G + T)$, and GC content were computed over the gene body on the gene strand, and genes were divided into fifths by each.

### Measured start sites in K562 and HCT116

The ENCODE plus- and minus-strand signals were read as integer counts of RNA 5′ ends (18.7 million reads). Strand labels were confirmed by checking that reads within 250 nt of annotated starts of protein-coding genes fell mostly on the gene strand (65.2% on chromosome 1). Sites were positions with at least 1, 3 or 10 reads at the eligible positions defined above, merged within 50 nt; sites with at least 10 reads are called strong. The activity of a gene was the number of reads on its strand within 250 nt of any of its annotated transcript starts: silent, 0; weakly active, 1–4; active, 5 or more, divided into thirds of genes. Strand ratios, deficits and confidence intervals were computed as for the predicted sites.

To identify silent genes reached by the transcription of neighbors, each active protein-coding or lncRNA gene was taken to cover its own strand from its start to 20 kb beyond its end, and the opposite strand for 5 kb upstream of its start. A silent gene was classified by whether any part of its body was covered on its gene strand, on its opposite strand, on both or on neither. The analysis was repeated with 50 kb and 10 kb, and with neighbors counted as active when they had at least one read at their start (Supplemental Table S1).

DNA methylation of a gene was the mean methylation of the CpGs at its eligible positions. Methylated and total reads were summed over the two strands of each CpG and over the two replicates, and CpGs with at least 10 reads were used (23.1 million; Pearson r of methylation between the replicates, 0.96); every CpG lay at a CG of the reference genome. Genes with fewer than 20 such CpGs were left out, which kept 9,176 of the 10,087 active protein-coding genes. Within each third of host activity, active genes were divided into thirds of methylation. Before seeing the data, we specified one comparison: the deficit of strong sites in the most minus the least methylated third, each third pooled over the three activity thirds, with genes resampled within each combination of activity and methylation third (2,000 resamples). After seeing the results, we added an equal-weight comparison, the mean of the differences within the three activity thirds. As a negative control, intergenic 1-Mb blocks were divided into thirds of their methylation, and their plus and minus strands were compared. The analysis was repeated with the earlier processing of the sequencing data.

HCT116 PRO-cap (experiment ENCSR165QKR) was processed by ENCODE in two ways from the same two replicates. Before seeing the data, we chose as primary the processing used in the default ENCODE analysis of the experiment (ENCAN786UJR): plus-strand files ENCFF440UVY and ENCFF078CJH and minus-strand files ENCFF103FBH and ENCFF907PJM, summed over the replicates (45.5 million autosomal reads, 2.4 times the K562 depth). The other processing (ENCFF266IRD, ENCFF209TJK, ENCFF116OZK and ENCFF828GNJ) served as a sensitivity analysis. Strong sites had at least max(3, round(10 × N / 18,699,448)) reads, N being the autosomal reads of the data set: 10 in K562 and 24 in HCT116. Eligible positions, sites and gene activity were defined as for K562, and the per-gene counts of the K562 analysis were reproduced exactly. H3K36me3 peaks were ENCODE replicated peaks (ENCFF520LHY for K562 and ENCFF029GQD for HCT116); they covered 0.6–9.3% of the eligible positions of active protein-coding genes. A site lay inside a peak when the position of the site with the most reads did, and the fifth of a site was set by the relative position of the site along the annotated gene, from the 5′ to the 3′ end. Before seeing the data, we recorded three primary comparisons in active protein-coding genes, each with the predicted sign: the deficit inside minus outside H3K36me3 peaks (both cell lines; positive), the deficit in the 3′ minus the 5′ fifth (both cell lines; positive) and the deficit in HCT116 (positive). Genes were resampled within the activity thirds (2,000 resamples), with two-sided bootstrap P values and no correction for multiple testing. Controls were silent genes and intergenic 1-Mb blocks (plus divided by minus strand); silent genes had too few strong sites (53–78 per strand) for the comparison of fifths. With the other HCT116 processing, the three HCT116 estimates were 15.2% (95% CI 8.6–21.4%), 6.3 percentage points (−98.8 to 48.2) and −41.0 (−66.4 to −19.4).

After seeing these results, we specified further analyses, each before it was run; they are exploratory. The distance of a strong site, or of the midpoint of a start-site pair, from the annotated 5′ end of the host gene was grouped into bins of 1–5, 5–10, 10–20, 20–40, 40–80 and at least 80 kb; because eligible positions lie at least 1 kb from annotated transcript starts, the first bin starts at about 1 kb. For strong sites, we also measured the distance from the nearest annotated transcript start of the same gene upstream of the site, which accounts for alternative promoters (within 5 kb, deficit 37%, 95% CI 29–45%, in K562 and 45%, 37–53%, in HCT116; beyond 5 kb, 2%, −7 to 9%, and −3%, −14 to 6%), and we checked that the deficit in the 5′ fifth held in all thirds of gene length. For the pairs, the host gene was found by the GENCODE identifier or, failing that, as the only protein-coding gene containing the midpoint of the pair on the strand of the sense start; all depth-matched pairs were placed, and the overall bias reproduced the main analysis. The bias of the pairs in a bin was computed as in the main analysis from the depth-matched pairs in that bin, with host genes resampled. A site or pair was near a CpG island when it lay within 2 kb of one. To separate the start of the gene from CpG islands, sites and pairs were divided into four groups (within 5 kb of the 5′ end of the host gene or farther, crossed with within 2 kb of an island or farther), and both cell lines were pooled, with genes resampled jointly for both lines, so that a gene present in both lines had the same weight in both (2,000 resamples). The readouts were the strand difference within 5 kb minus beyond 5 kb, both more than 2 kb from any island, and the strand difference beyond 5 kb near an island minus away from islands. Finally, the excess within 5 kb (within minus beyond) was compared between the most and the least active third of genes: for sites, the activity thirds defined above; for pairs, thirds of the reads at the annotated start of the host gene, over the host genes of the depth-matched pairs in each line. The 5-kb boundary was the upper end of the first distance bin. For Figure 3F, drawn after these analyses, strong sites were counted in 1-kb bins of distance from the 5′ end of the gene up to 30 kb (sites farther away, about half of all, are included in Supplemental Fig. S5B); the strand ratio was computed in 3-kb windows centered on each bin, with 95% CIs from 2,000 resamples of genes within thirds of gene activity, and the density as sites per 1,000 active genes whose annotated span reaches the bin. All estimates are in Supplemental Fig. S5, Figure 3F and the Source Data of these figures.

To test the deficit and its position with a second assay, we used the K562 GRO-cap library with TAP of Core et al. (2014) (GEO GSM1480321, series GSE60456; the library without TAP, GSM1480322, was not used), recorded the questions before the analysis, and ran the scripts of the PRO-cap analysis unchanged. The authors' files hold single-base counts of RNA 5′ ends on hg19 (26.1 million autosomal reads). We transferred each position to hg38 through the ungapped blocks of the UCSC hg19ToHg38 chain: positions in inverted blocks changed strand (112,520 reads), positions outside any block were dropped (15,175 reads), and so were reads that landed on sequences other than chromosomes 1–22, X, Y and M (3,650 reads), which kept 99.93% of the reads. The transfer agreed with an independent position-by-position mapping on synthetic data. In hg38, 58.3% of the reads within 250 nt of annotated protein-coding starts on chromosome 1 lay on the gene strand, so the strand labels were used as given. Strong sites had at least 14 reads (the K562 threshold scaled to 26.0 million autosomal reads). The three predictions, each tested in active protein-coding genes with genes resampled within activity thirds (2,000 resamples, two-sided bootstrap P, no correction), were a deficit above zero, a smaller deficit in the 3′ than in the 5′ fifth, and a larger deficit within the first 5 kb (the first distance bin) than beyond. In secondary analyses, the deficit rose with activity (5%, 11% and 19% from the least to the most active third), and the deficit inside minus outside K562 H3K36me3 peaks was 22.2 percentage points (95% CI −0.3 to 39.8, P = 0.053), opposite in sign to the PRO-cap estimate (−18.1, −73.3 to 18.0).

### Population test

All protein-coding intronic variants, and random subsets of lncRNA intronic (30%) and intergenic (20%) variants at eligible positions, were scored with Puffin-D, with the reference and with the alternative allele, reading the prediction within 50 nt of the variant on each strand. To save computing time, variants at least 1,000 nt apart were first scored together in shared 100-kb windows, and every variant whose effect in this screen exceeded 0.003, or that came within 0.003 of a site threshold, was scored again on its own; in a random audit of the remaining variants, 0.13–0.49% of the audited variants, depending on the chromosome, had an effect above 0.003 that the screen had missed. Only sites where the number of genotyped alleles was at least 80% of the maximum (152,430) were used, which left 215.5 million variants in the three sets.

With Puffin-D, variants were classified as follows. A variant created a start when, on one strand, the reference prediction was below the weak (0.0439) or medium (0.1432) threshold and the alternative was at or above it; it removed a start in the reverse case. It strongly raised a start when the change was at least 0.01 on one strand and below 0.003 on the other (a fall on the other strand was allowed). It moderately raised a start when the change was at least 0.003 but below 0.01 on one strand and smaller than 0.003 in absolute value on the other. It strongly lowered a start when the change was −0.01 or less on one strand and smaller than 0.003 in absolute value on the other. It was neutral when the change was below 0.001 in absolute value on both strands. The expected share of singletons (allele count 1) for each class was the mean, over its variants, of the share of singletons among neutral variants of the same set with the same three-base context and alternative base, oriented to the gene strand (192 contexts). The excess of singletons was the observed share minus this expectation (Lek et al. 2016). The analysis was repeated without CpG sites, with contexts further split into five bins of local GC content (±50 nt), and with both. Positive controls were variants at canonical splice sites (the first two and last two bases of introns of canonical protein-coding transcripts) and at positions 3–6 of splice donors. As a check of calibration, the expectation from neutral variants in half of the genes was applied to neutral variants in the other half (excess −0.0008). Differences between classes were bootstrapped over genes (1-Mb blocks for intergenic DNA), resampling the same units for both members of each difference (1,000 resamples).

The same variants were also scored with ProCapNet. Each variant was read with the model whose test group included its chromosome, in the 1,000-nt output tile of the genome-wide ProCapNet maps when the 101-nt readout fitted inside it, and otherwise in a tile shifted by 500 nt; the reference and the alternative were read from the same window, and every alternative allele was scored on its own, without a screen. For variants in unshifted tiles the reference readout reproduced the genome-wide tracks on every chromosome (99th percentile of the relative difference below 0.5%, the precision at which the tracks are stored). Site thresholds were those of the ProCapNet maps (1.131 and 2.658). Because ProCapNet predicts counts rather than Puffin-D's scaled signal, its cut-offs for no effect (0.215), for moderate change and for the other strand (0.495) and for strong change (1.091) were set so that the same share of strand changes passed them as passed Puffin-D's cut-offs on chromosomes 8 and 9 (5.3%, 1.7% and 0.48% of changes). The test was otherwise identical. All classes, sets and settings are in Supplemental Table S1.

### Statistical analyses

Confidence intervals are 95% percentile bootstrap intervals; the resampled units (host genes, genes, 1-Mb blocks) and the number of resamples are given in each section and figure legend. Random seeds were fixed (seed 1; seed 1001 for the analyses resampling host genes in Figs. 1B and 2A). Tests are two-sided. The number of units (pairs, genes, blocks or variants) is given in each figure legend. Conclusions rest on the confidence intervals of a small number of primary comparisons: the depth-matched asymmetry, the share predicted by held-out ProCapNet models, the gene-strand deficit of potential and measured start sites, and the sense-minus-antisense difference in the excess of singletons. For DNA methylation, one comparison was specified before the data were seen, and the comparison added afterwards is labeled as such. For strong measured sites, three comparisons were specified before the data were seen (the deficit in HCT116, inside minus outside H3K36me3 peaks, and in the 3′ minus the 5′ fifth of genes); the analyses of distance from the start of genes, of CpG islands and of activity near the start were specified after these results, each before it was run, and are labeled exploratory. The repetition with K562 GRO-cap data (the deficit, the 3′ minus the 5′ fifth and the first 5 kb minus beyond) was specified before those data were analyzed. P values are reported without correction for multiple testing, and the robustness analyses (depth bands, distance to annotated starts, CpG and GC matching, other neighbor zones) are presented as consistency checks rather than as independent tests.

### Software

All analyses were run with scripts written for this study, numbered by analysis step (02–29); they are archived, together with the result tables, in a versioned release at Zenodo (https://doi.org/10.5281/zenodo.24161109) and are available at https://github.com/michal-szczesniak/intronic-start-asymmetry. Analyses used Python 3.11.17 with NumPy 2.4.6, pandas 3.0.6, SciPy 1.17.1, pyfaidx 0.9.0.4, pyBigWig 0.3.26 and PyTorch 2.3.1 (CUDA 12.1); figures were drawn with matplotlib 3.11.2 directly from the result tables. The full software environment is listed in env/purge.yml of the archived code. An AI assistant (Claude, Anthropic) was used to draft analysis code and text, which the authors checked and revised; every number was traced to its result file.

### Supplemental Figures

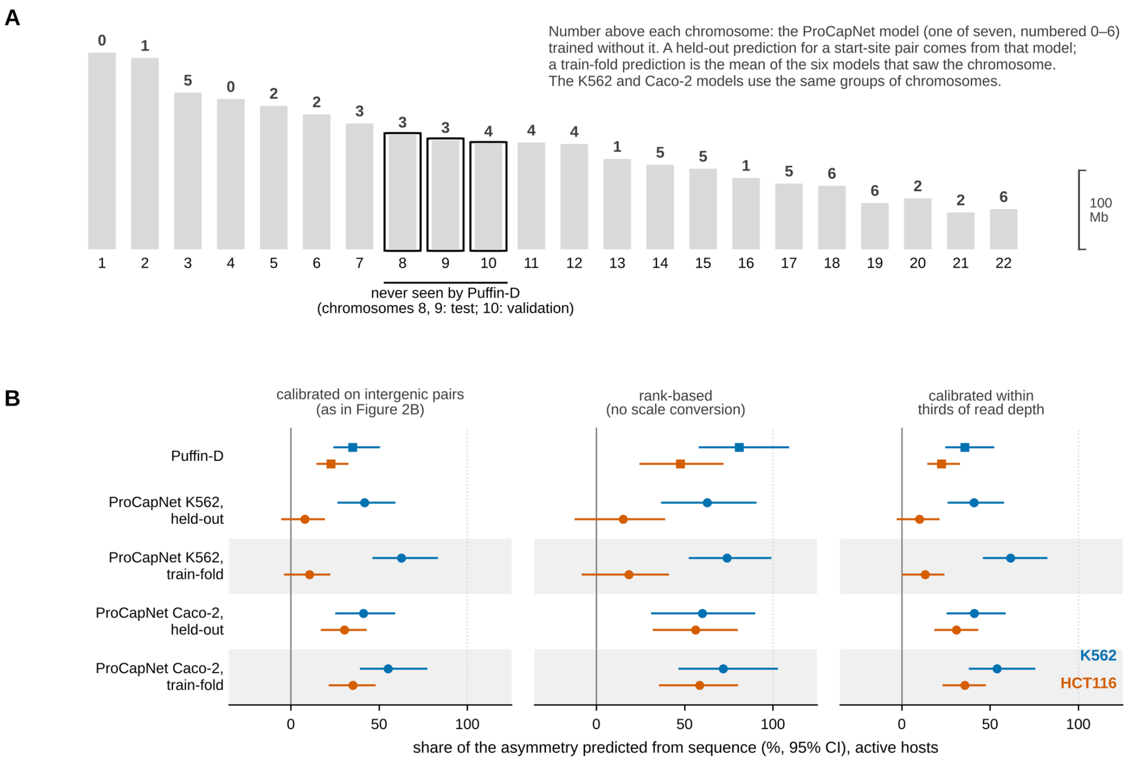


**Supplemental Figure S1. Which model saw which chromosome, and the share of the asymmetry predicted under three calibrations.** (A) Autosomes drawn to scale. The number above each chromosome is the ProCapNet model (one of seven, numbered 0–6) trained without it; the K562 and Caco-2 models use the same groups of chromosomes. Boxed, chromosomes never seen by Puffin-D (8 and 9, test; 10, validation). (B) Share of the asymmetry of intronic pairs in active hosts that is predicted from sequence, in K562 (blue) and HCT116 (orange-red), by Puffin-D (squares) and ProCapNet (circles). Held-out, the ProCapNet model that never saw the pair's chromosome; train-fold (shaded rows), mean of the six models that saw it. Left, calibration by least squares through the origin on intergenic pairs taken in both orientations (as in Fig. 2B); middle, rank-based share (area under the curve minus 0.5, predicted divided by measured), which needs no calibration; right, calibration within three bands of read depth. Bars, 95% CI from 2,000 resamples of host genes, with intergenic pairs resampled in 1-Mb blocks. n = 1,611 (K562) and 2,489 (HCT116) intronic pairs; 3,933 and 6,193 intergenic pairs.

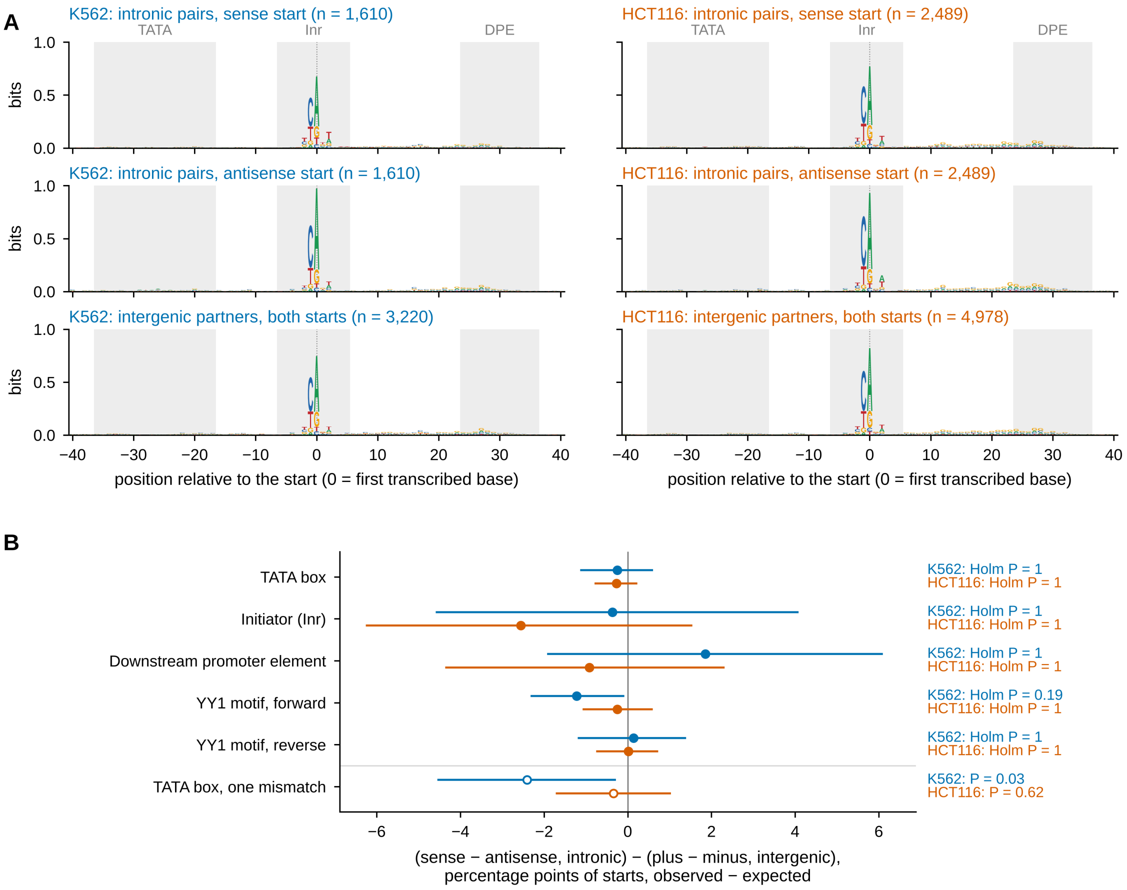


**Supplemental Figure S2. Known core promoter elements do not distinguish the two starts of intronic pairs.** Sequence around the two starts of the depth-matched pairs (K562, blue: 1,610 pairs in 964 host genes; HCT116, orange: 2,489 pairs in 1,187 host genes). (A) Sequence logos (information content in bits) from 40 nt before to 40 nt after each start (position 0, first transcribed base), for the sense and the antisense start of intronic pairs and for both starts of their intergenic partners. Gray: positions that the TATA box, the initiator (Inr) and the downstream promoter element (DPE) may cover. (B) For each element, the share of starts that carry it at its usual position, observed minus expected from 20 dinucleotide shuffles of the surrounding 121 nt, as the difference between the sense and the antisense start of intronic pairs minus the difference between the plus and the minus start of their intergenic partners, in percentage points of starts; 95% CIs from 2,000 bootstrap resamples of host genes. Filled symbols: the five elements chosen before the analysis, with Holm-corrected two-sided P over these five within each cell line; open symbols: TATA box with at most one mismatch (sensitivity analysis, uncorrected P). Definitions are given in the Supplemental Methods.

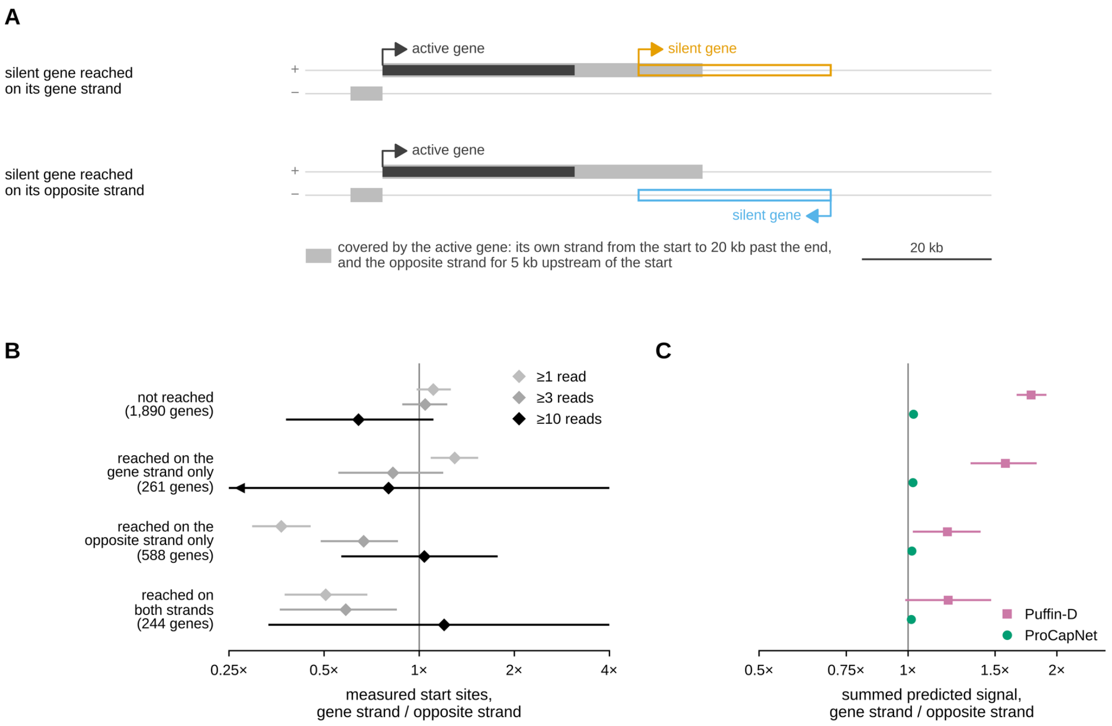


**Supplemental Figure S3. In silent genes, the strand difference of weak measured sites follows the strand reached by transcription from active neighbors.** (A) The rule, drawn to scale: each active protein-coding or lncRNA gene covers its own strand from its start to 20 kb past its end, and the opposite strand for 5 kb upstream of its start. A silent gene (no PRO-cap reads within 250 nt of any of its annotated starts) is classified by whether any part of its body is covered on its gene strand, on its opposite strand, on both or on neither. (B) Measured start sites in K562 with at least 1, 3 or 10 reads, gene strand divided by opposite strand, in introns of silent protein-coding genes of each class. (C) Summed predicted signal, gene strand divided by opposite strand, for the same genes: Puffin-D (squares) and ProCapNet (circles). Strand ratios on a logarithmic axis; bars, 95% CI from 2,000 bootstrap resamples of genes; arrowhead, interval continues beyond the axis. Genes per class are given in B (2,983 silent genes in all; one silent gene is missing from the Puffin-D tables).

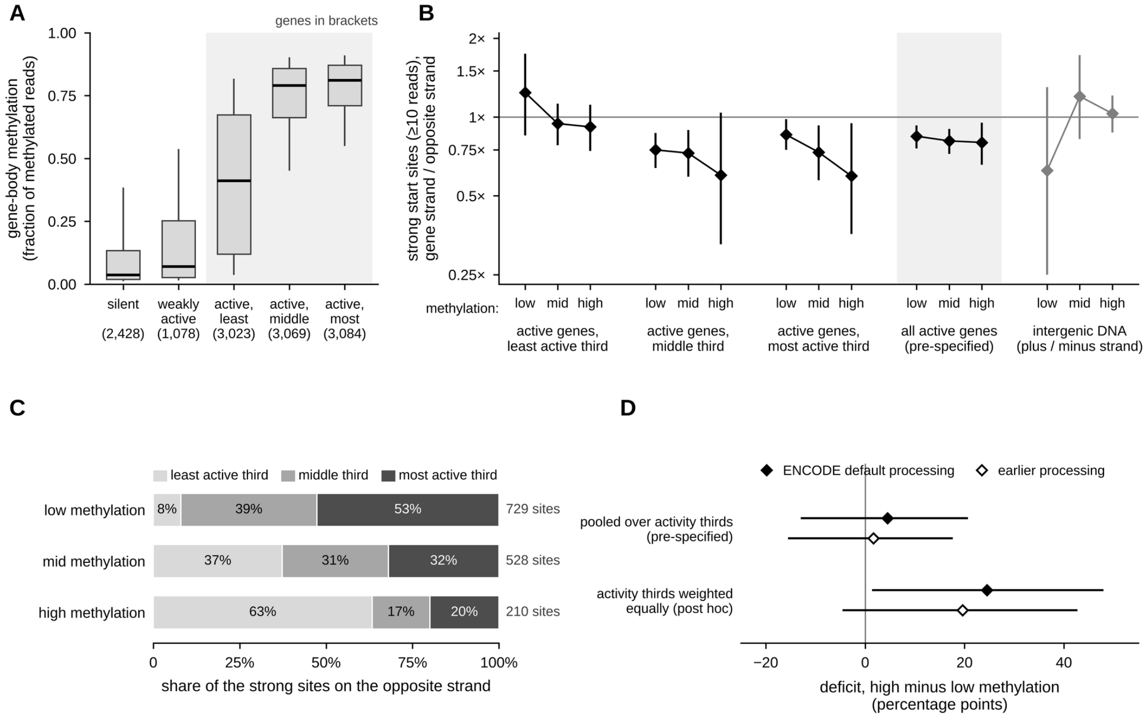


**Supplemental Figure S4. Gene-body DNA methylation and strong measured start sites in K562.** Methylation from ENCODE whole-genome bisulfite sequencing (ENCSR765JPC): for each protein-coding gene, the mean methylation of the CpGs with at least 10 reads at its analyzed intronic positions; genes with at least 20 such CpGs (9,176 of 10,087 active genes). (A) Methylation by host activity, groups as in Figure 3C (number of genes in brackets); boxes, quartiles; thick line, median; whiskers, 10th and 90th percentiles. (B) Strong measured sites (at least 10 reads), gene strand divided by opposite strand, in thirds of methylation (low, mid, high) formed within each third of active genes (1,007–1,028 genes per point; median methylation 0.07, 0.41 and 0.75 in the least active third, and 0.58–0.89 in the other two); the same thirds pooled over the three activity thirds (shaded; the pre-specified comparison; 3,058–3,059 genes per point; deficits 16%, 95% CI 7–24%; 19%, 10–28%; and 20%, 5–34%); and, as a negative control, intergenic 1-Mb blocks in thirds of their methylation (plus strand divided by minus strand; 848–849 blocks per point). Values below 1 mean fewer sites on the gene strand. (C)* *Where the strong sites of each methylation group come from: the share of its strong sites on the opposite strand contributed by each third of active genes (number of sites at right). The planned comparison adds up the sites of the three activity thirds, so each activity third counts in proportion to its share. Among highly methylated genes, most sites (63%) come from the least active third, which has no detectable deficit; among genes of low methylation, only 8% do. (D) Difference in the deficit between high and low methylation, in percentage points (positive: fewer sites on the gene strand in highly methylated genes). Top, the planned comparison, which adds up the sites of the three activity thirds (4.5, 95% CI −12.9 to 20.7, P = 0.61). Bottom, the average of the three differences computed within each activity third, which gives each activity third the same weight (chosen after seeing the data; 24.5, 1.4 to 48.0, P = 0.04). Filled symbols, methylation from ENCODE's current default processing (files ENCFF660IHA and ENCFF328NMN); open symbols, from an older ENCODE processing of the same sequencing reads (files ENCFF867JRG and ENCFF721JMB; 1.7, −15.5 to 17.6, and 19.6, −4.6 to 42.7), shown to check that the result does not depend on how the reads were processed. In the least active third, methylation also rose with host activity (Spearman $\rho = 0.54$), so part of its difference may reflect activity. Bars, 95% CI from 2,000 bootstrap resamples of genes within each combination of activity and methylation third (intergenic: 1-Mb blocks); P, two-sided bootstrap. All values are in Supplemental Table S1 (sheet S1H).

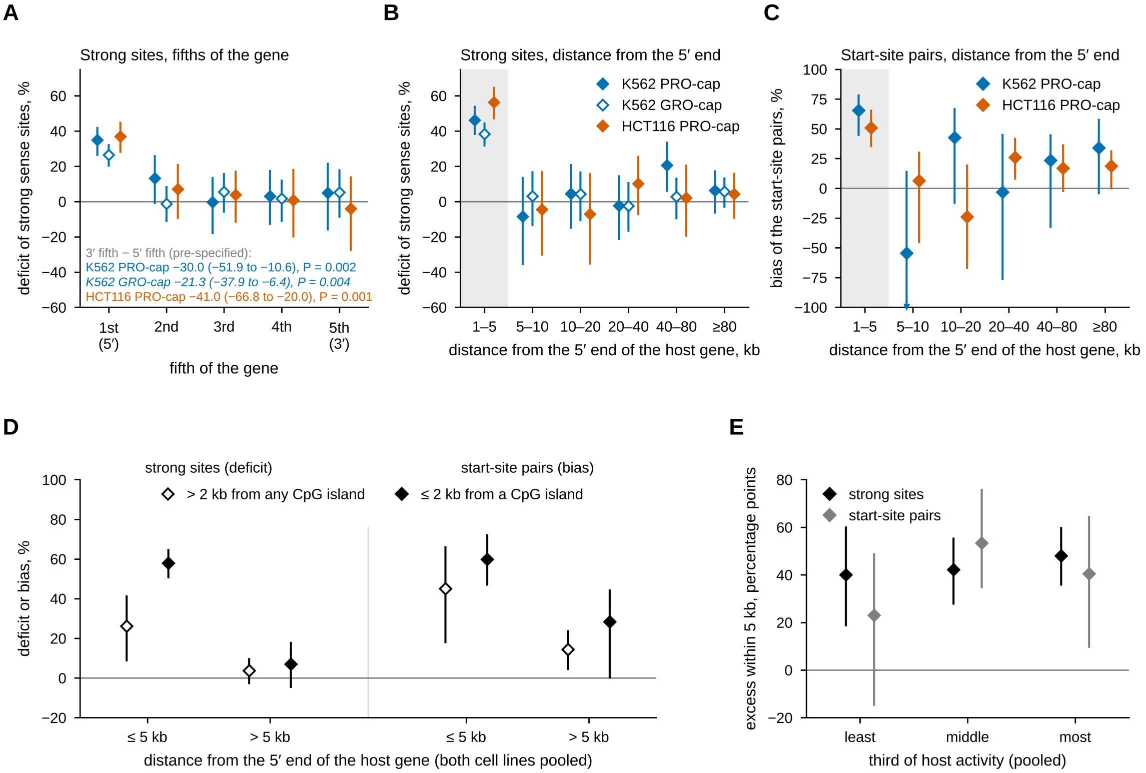


**Supplemental Figure S5. The gene-strand deficit of strong measured sites lies near the start of genes.** Strong measured start sites in active protein-coding genes (K562 PRO-cap, filled blue: at least 10 reads, 10,087 genes; HCT116 PRO-cap, orange: at least 24 reads, 10,538 genes; in A and B also K562 GRO-cap, open blue: at least 14 reads, 9,907 genes) and depth-matched start-site pairs (1,610 in K562 and 2,489 in HCT116). Deficit, 100 × (1 − sense / antisense strong sites); bias of the pairs, 100 × (1 − 2^d), where d is the median log2 ratio of the intronic pairs minus that of their depth-matched intergenic partners. Positive values mean fewer or weaker starts on the gene strand. (A) Deficit in fifths of the annotated gene, from the 5′ to the 3′ end; the contrast 3′ fifth minus 5′ fifth was specified before the data were seen (two-sided bootstrap P). (B) Deficit by distance of the site from the annotated 5′ end of the host gene; eligible positions lie at least 1 kb from annotated transcript starts, so the first bin (gray) starts at about 1 kb; the same sites in 1-kb bins up to 30 kb are shown in Figure 3F. (C) Bias of the pairs by distance of the pair from the 5′ end of the host gene. (D) Both cell lines pooled: deficit and bias within 5 kb of the 5′ end or farther, for sites and pairs more than 2 kb from any CpG island (open symbols) or within 2 kb of one (filled symbols). (E) Both cell lines pooled: excess within 5 kb (deficit or bias within 5 kb minus beyond 5 kb, in percentage points) in thirds of host activity. Panels B–E show exploratory analyses chosen after the PRO-cap results in A; the GRO-cap estimates in A and B were tested with predictions recorded in advance. Bars, 95% CI from 2,000 bootstrap resamples of genes (host genes for pairs; in D and E, genes drawn jointly for both cell lines); arrowhead, interval continues beyond the axis. Numbers of sites and pairs in each group are given in the Source Data.

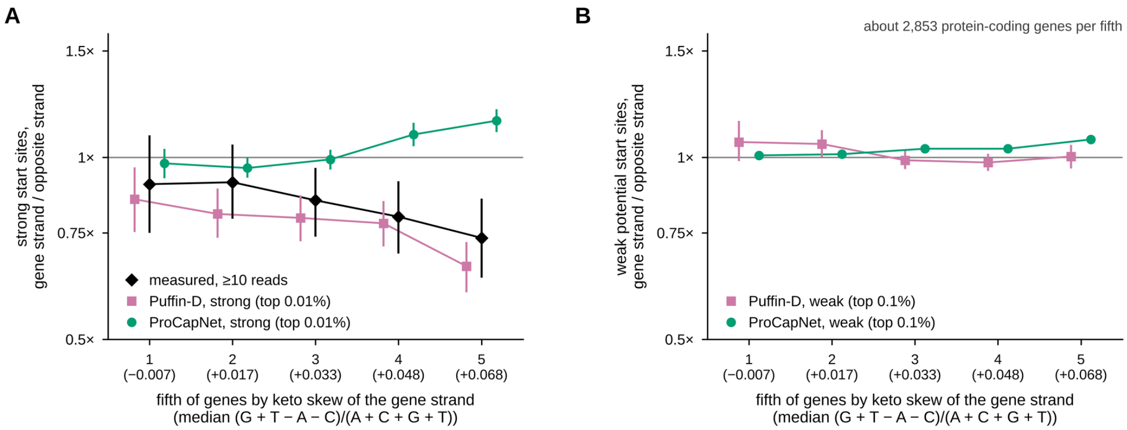


**Supplemental Figure S6. Strand ratios of strong and weak start sites by the keto skew of the gene strand.** Protein-coding genes were divided into fifths by the keto skew of the gene body on the gene strand, (G + T − A − C)/(A + C + G + T); the median of each fifth is given in brackets (2,852–2,854 genes per fifth). (A) Strong sites: measured sites in K562 with at least 10 reads (diamonds), and potential sites among the highest 0.01% of predictions of Puffin-D (squares) and ProCapNet (circles), gene strand divided by opposite strand. (B) Weak potential sites (highest 0.1% of predictions). Strand ratios on a logarithmic axis; bars, 95% CI from 2,000 bootstrap resamples of genes. Change in the deficit of strong sites from the lowest to the highest fifth (percentage points, 95% CI): measured, 16.7 (−2.6 to 37.4); Puffin-D, 19.2 (6.9 to 31.4); ProCapNet, −17.2 (−24.8 to −9.7).

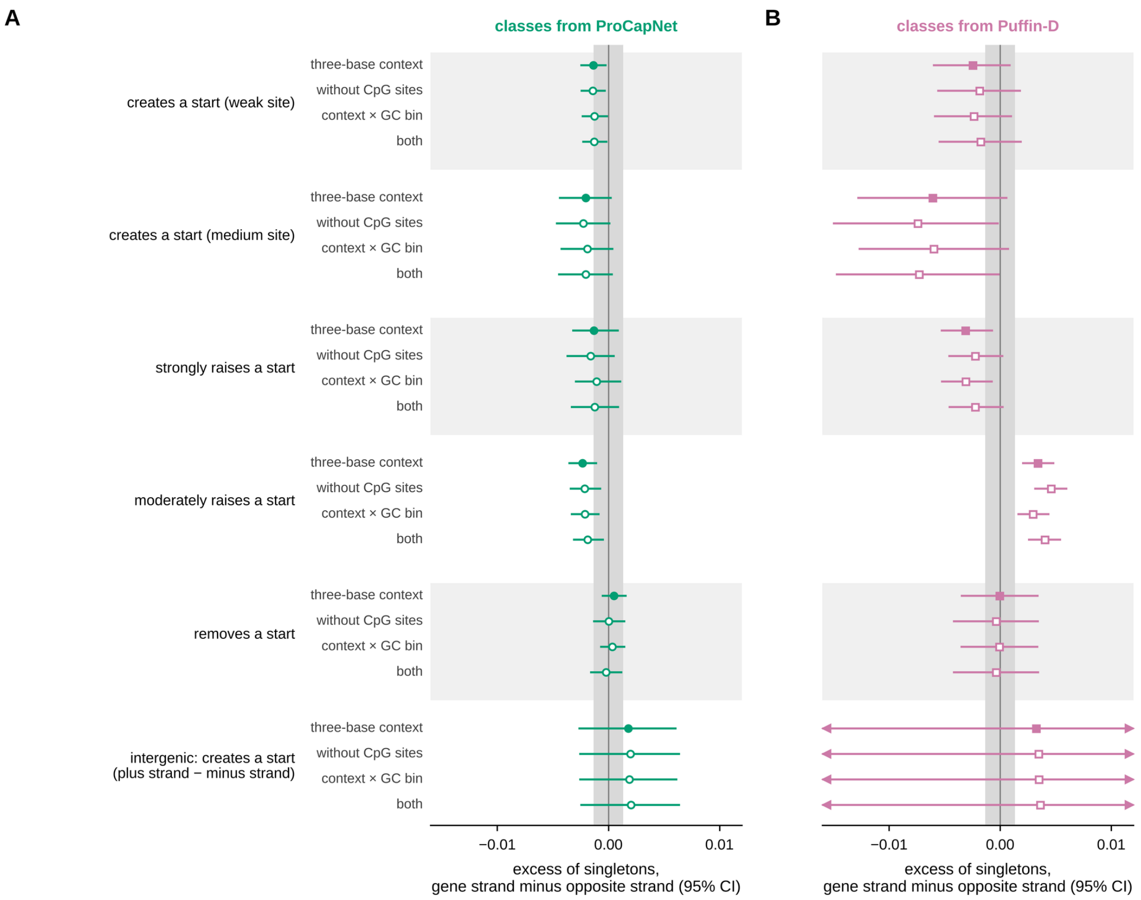


**Supplemental Figure S7. The strand comparison of the population test under four definitions of the expectation.** Gene-strand minus opposite-strand excess of singletons (intergenic: plus minus minus strand) for classes from ProCapNet (A) and Puffin-D (B). The expectation came from variants of the same mutation type and three-base context that have no predicted effect (filled symbols; the main analysis, Fig. 4C), from the same without CpG sites, with contexts further split into five bins of local GC content (±50 nt), or with both. Gray band, ±1% of the excess of singletons at canonical splice sites (one hundredth of that value); bars, 95% CI from 1,000 bootstrap resamples of genes (intergenic: 1-Mb blocks); arrowheads, interval continues beyond the axis. Open symbols, the three alternative expectations.

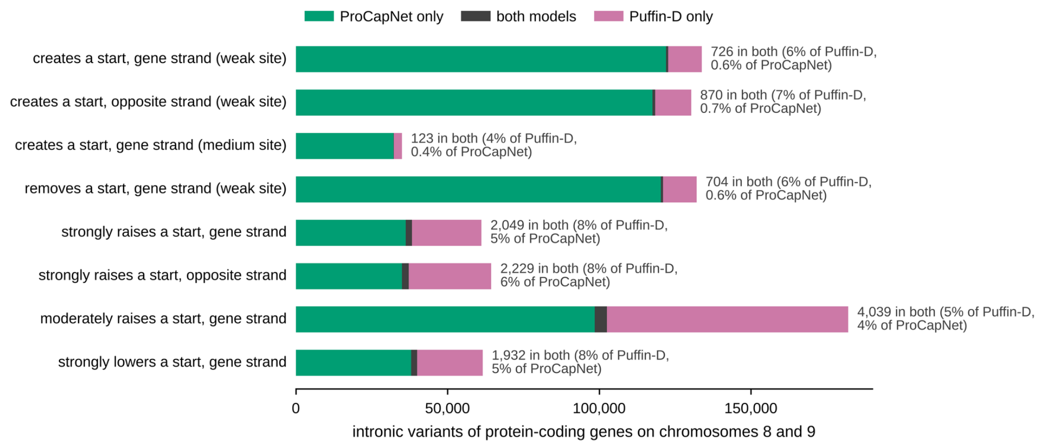


**Supplemental Figure S8. ProCapNet and Puffin-D classify largely different variants.** Intronic variants of protein-coding genes on chromosomes 8 and 9 in each class of predicted effect, classified by ProCapNet only (green), by both models (dark gray) or by Puffin-D only (pink); at right, the shared variants as a percentage of each model's class. Weak and medium sites as in Figure 3B; cut-offs for strong and moderate changes matched between the models (Supplemental Methods). Variants classified as having no effect: ProCapNet, 16,001,124; Puffin-D, 16,012,573; both, 14,709,254.
